# The dynamic and heterogeneous structure of the non-canonical inflammasome

**DOI:** 10.64898/2026.04.15.718739

**Authors:** Alexander I.M. Sever, James M. Aramini, Jeffrey P. Bonin, Huaying Zhao, Hanlin Wang, John L. Rubinstein, Peter Schuck, Lewis E. Kay

## Abstract

Inflammasomes are high molecular weight complexes that play an integral role in the innate immune system, triggering an inflammatory cascade to protect against cellular stresses such as pathogenic bacteria. Both canonical and non-canonical inflammasomes have been described in the literature and detailed structural studies of many components of the more complex and larger canonical versions have been reported. In contrast, corresponding structures of the non-canonical inflammasome have not emerged even though it consists of only two components: lipopolysaccharide (LPS) from gram-negative bacteria, and one of caspase-4 or caspase-5 in humans or caspase-11 in mice. Here we determine the stoichiometry of the non-canonical inflammasome using size-exclusion chromatography coupled with UV, refractive index, and light-scattering measurements, showing that the non-canonical inflammasome is heterogeneous, comprised of three major complexes with different numbers of LPS and caspase molecules. Solution Nuclear Magnetic Resonance (NMR) spectroscopy studies of the N-terminal Caspase Activation and Recruitment Domain (CARD) of caspase-11, that binds LPS, show that it is largely unstructured in the absence of lipid, with pervasive dynamics on the μs-ms timescale. Formation of this complex increases the alpha-helical content of the CARD but the dynamics persist, multiple conformers are formed, and tertiary contacts are transient, consistent with formation of a molten globule. Our NMR results establish that the protease domain of caspase-11 is monomeric in isolation. As proteolysis is linked with dimerization, the protease domains are inactive in this state, but upon formation of the non-canonical inflammasome dimerization occurs, priming the complex for rapid processing of substrates.

**Significance Statement:** Animals respond to injury, infection, or toxic materials via a process called inflammation. At the molecular level inflammation involves formation of large machines - inflammasomes - that are instrumental in triggering cascades that lead to an immune response. Although structural studies of canonical inflammasomes have emerged, much less is known about the structures of non-canonical inflammasomes. Using solution Nuclear Magnetic Resonance (NMR) spectroscopy in concert with other biophysical approaches we show that non-canonical inflammasomes are highly dynamic and structurally heterogeneous, and we characterize the different sized inflammasome particles that are formed in terms of their composition. We also show that once formed, non-canonical inflammasomes are primed to rapidly cleave substrate molecules, necessary for propagating the immune response.

## Introduction

Programmed cell death (PCD) pathways are essential for the development and survival of multicellular eukaryotes (1). Numerous forms of PCD have been discovered, with each having a unique trigger and function within the host. Apoptosis is a highly regulated form of PCD that is inherent to every cell within an organism and can be initiated upon cellular stresses. These include intrinsic cellular insults, such as irreparable DNA damage, triggering apoptosis through the formation of large protein complexes, called apoptosomes, which activate caspase-9 (Casp9) proteases via molecular crowding on the apoptosome surface. Detailed cryo-EM studies have produced high resolution images of the apoptosome scaffold (2–4), while solution Nuclear Magnetic Resonance (NMR) studies have added to these results by focusing on the protease domains of Casp9 in the context of the apoptosome, as these are not visible in structures of the complex (5).

An analogous PCD pathway, known as pyroptosis, is utilized exclusively by innate immune cells to trigger an inflammatory response in the host (*SI Appendix*, **Fig. S1**). This cascade is initiated during an infection when evolutionarily conserved microbial components, known as pathogen-associated molecular patterns (PAMPs), are detected by host immune cells either extra-or intra-cellularly via interaction with a diverse class of proteins called pattern recognition receptors (PRRs) (6). A subset of these PRRs, called canonical inflammasomes, are responsible for the formation of megadalton-sized complexes in response to PAMPs (7). These large, multi-component machines form filamentous structures that bind and activate Casp1, which can then cleave pro-inflammatory cytokines, such as pro–interleukin-1β (pro-IL-1β) and pro–IL-18 (8–10). Additionally, the pore-forming protein gasdermin D (GSDMD) is also activated during this process (11–14). It embeds in cell membranes to enable the escape of mature interleukins to the extracellular matrix (15–17) where they can bind to surface receptors on neighbouring cells, propagating the inflammatory response. As with the apoptosome, detailed X-ray and cryo-EM studies have provided insights into the molecular architectures of the different variants of canonical inflammasomes (18–23).

The inflammatory cascade can also proceed through the formation of simpler structures, termed non-canonical inflammasomes (24, 25) (*SI Appendix*, **Fig. S1**). These are formed through the direct association of Casp4 or Casp5 in humans or Casp11 in mice with cytosolic lipopolysaccharide (LPS), initiating pyroptotic cell death (26–30). LPS, considered the prototypical PAMP, is a major component of the outer membrane of gram-negative bacteria, and is composed of Lipid A, the core sugars, and the O-antigen. While the core sugars and O-antigen have significant variability amongst species, the Lipid A moiety remains highly conserved, providing an ideal ‘pattern’ to target (31). Mouse Casp11 (Uniprot: P70343, referred to as Casp11 in what follows) consists of 373-residues and contains two functional domains connected via a long, disordered linker (**Fig. 1A**; top). The N-terminal Caspase Activation and Recruitment Domain (CARD; residues 1 to 80) is responsible for interacting with the Lipid A moiety of LPS (29, 32), while the C-terminal Protease Domain (PD, residues 102-373) exhibits peptidase activity via a catalytic cysteine residue (Cys 254) (29, 33). The PD can be further subdivided into p20 (residues 102-256) and p10 (residues 292-373) domains, which are connected via another disordered linker that contains an auto-cleavage site (Asp 285) (34–36). Like other initiator or inflammatory caspases, the Casp11 PD is a monomer in its resting state with dimerization required for proteolytic activity. Whether the effective concentration of Casp11 in the non-canonical inflammasome is sufficient to support dimerization of PDs or whether dimerization is triggered through substrate binding, as for Casp9 (5), has yet to be established (**Fig. 1A**; bottom).

**Figure 1.**
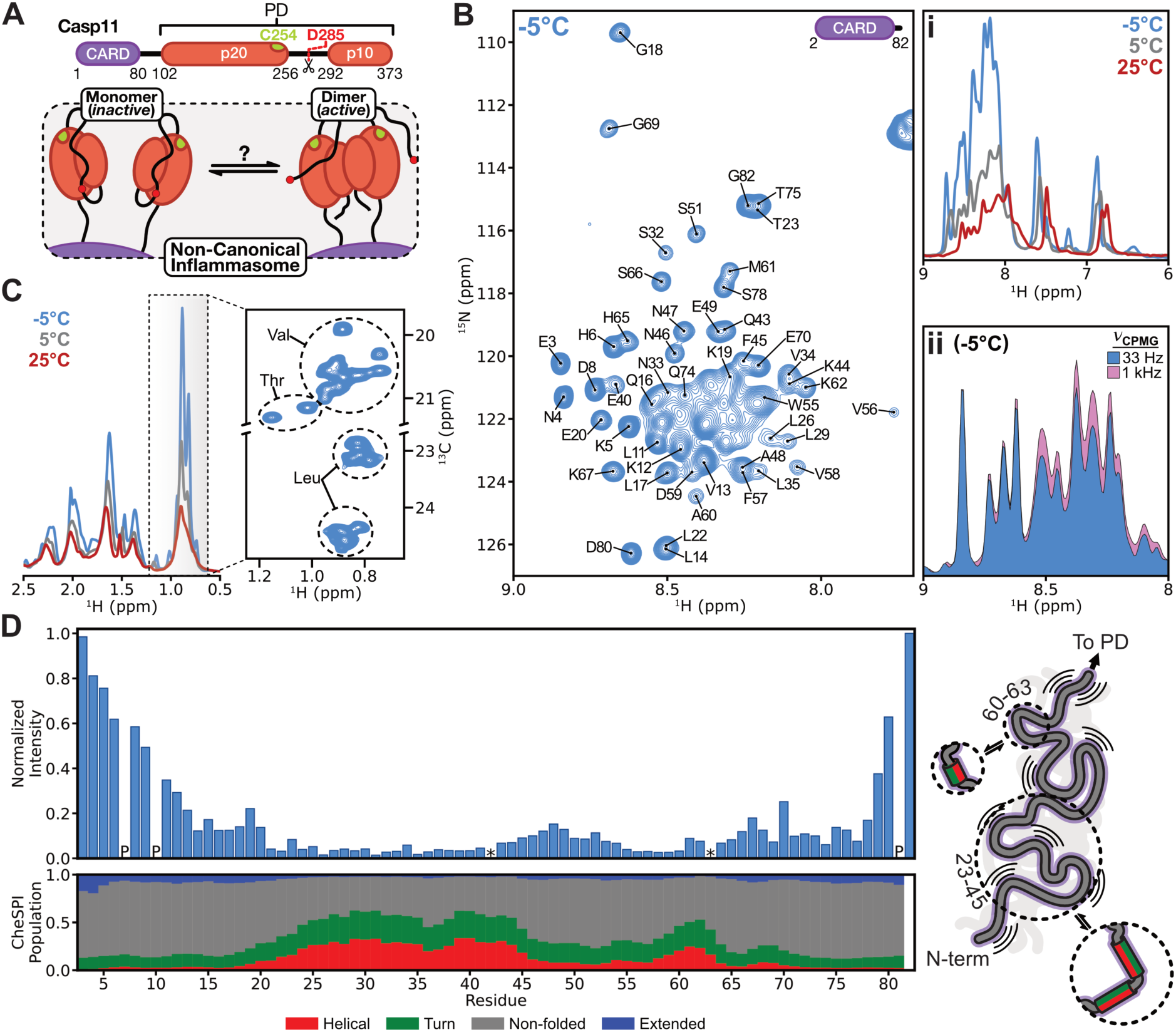
(A) (top) Domain organization of mouse caspase-11 (Casp11) with residue numbers at the start and end of the folded domains. The catalytic cysteine (Cys 254; green sphere) and an auto-cleavage site in the p20-p10 linker (Asp 285; red dotted line) are illustrated. (bottom) Activation of the non-canonical inflammasome is accompanied by dimerization and subsequent auto-cleavage at Asp 285 (34–36). The dimerization status of the PDs is the subject of the current study. **(B)** (left) 2D ^1^H-^15^N HSQC spectrum of U-^13^C,^15^N Casp11 CARD recorded at 14.1 T (600 MHz, ^1^H frequency) and -5°C. Assignments of some resonances are indicated (see *SI Appendix*, **Fig. S1A** for complete annotations). (i, top right) 1D ^15^N-edited ^1^H spectra of U-^13^C,^15^N Casp11 CARD recorded at either 25°C (red trace), 5°C (grey trace), or -5°C (blue trace) at 11.7 T (500 MHz, ^1^H frequency). (ii, bottom right) 1D ^1^H-detected ^15^N CPMG-based spectra of U-^15^N Casp11 CARD recorded at low (33 Hz; blue-filled) and high (1 kHz; pink-filled) ϖ_CPMG_ values, 18.8 T (800 MHz, ^1^H frequency) and -5°C. **(C)** 1D ^13^C-edited ^1^H spectra of U-^13^C,^15^N Casp11 CARD, focused on the methyl region, and recorded as a function of temperature (left); the methyl-containing region of the 2D ^1^H-^13^C HSQC spectrum recorded at -5°C is shown on the right. Data acquired at 18.8 T (800 MHz, ^1^H frequency). **(D)** (top) Normalized intensity values of each residue from a 3D HNCO experiment. ‘P’ = proline and ‘*’ indicates significant overlap that prevented accurate quantification. (bottom) Secondary structure population estimates based on the CheSPI program (51), identifying regions with significant helical/turn propensities. A cartoon highlighting the highly dynamic nature of the Casp11 CARD is illustrated on the right. Regions with a helix + turn population > 40% (arbitrarily chosen), are annotated with dotted circles, highlighting the interconversion between unfolded (grey line) and helical/turn (red/green cylinders) segments.

Although a direct interaction between the Casp11 CARD and LPS was established over a decade ago (29), and shown to lead to proteolytic upregulation, structures of the intact non-canonical complex have not emerged. X-ray and AlphaFold models of the isolated CARD indicate that it adopts a helical bundle (37–39), but recent circular dichroism (CD) and hydrogen-deuterium exchange mass spectrometry (HDX-MS) studies challenge the validity of these structures, finding instead that the CARD is disordered and undergoes a structural transition upon interacting with LPS (40). Whether this structural transition results in stabilization of tertiary contacts, and if so whether these interactions reflect those seen in the X-ray/AlphaFold structures, is still unknown. Furthermore, the overall quaternary structure of the non-canonical inflammasome is contentious, with two models being proposed. In one case, it has been postulated that the CARD embeds directly into LPS micelles (41) at areas of positive membrane curvature without changes to the micelle structure (42). Alternatively, other studies suggest that the CARD disaggregates LPS micelles into smaller complexes, although the exact oligomeric state and CARD:LPS stoichiometry in this case remains unclear (32, 40). Finally, while high resolution structures of the Casp11 PD have been solved by X-ray methods (43), the PD dimerization affinity is not known, and as mentioned above, the oligomeric state of the PDs upon formation of the non-canonical inflammasome complex remains to be established.

Herein, using solution state Nuclear Magnetic Resonance (NMR) spectroscopy, we demonstrate that although the Casp11 CARD exists in a largely disordered state prior to LPS binding, specific regions of the domain form transient secondary structure and intramolecular contacts on the μs-ms timescale. Upon binding LPS, the helical content of the CARD domain increases, yet it remains highly dynamic, with sidechains adopting numerous conformations, reminiscent of a molten globule. Characterization of the oligomerization state of the non-canonical inflammasome complex via size-exclusion chromatography coupled to UltraViolet, Refractive Index and Light Scattering (SEC-UV-RI-LS) detectors reveals that under our experimental conditions, three distinct complexes consisting of four, six, and eight Casp11 molecules are formed. NMR studies quantifying the effective concentration of PDs in non-canonical inflammasomes indicate that the Casp11 PDs readily dimerize in this environment, priming the complex for rapid processing of its substrates. Together, these results highlight the dynamic and heterogeneous nature of the non-canonical inflammasome.

## Results

### The Casp11 CARD is highly dynamic

NMR spectroscopy is a powerful technique for identifying short-lived interactions and the presence of residual structure in dynamic systems (44, 45). In order to establish whether the CARD adopts a four-helix bundle in solution, as suggested by earlier X-ray studies (37), we recorded a ^1^H-^15^N heteronuclear single quantum correlation (HSQC) spectrum of a U-^13^C,^15^N labelled Casp11 CARD (residues 2-82) at 25°C and pH 6. The resulting spectrum was of poor quality and contained noticeably fewer than the expected number of peaks, with the majority being low intensity and broadened. While increasing the temperature or the pH (from 6 to 7.4) did not improve spectral quality, decreasing the temperature to -5°C led to a considerable increase in both the number of peaks and their intensities (**Fig. 1B**, **Fig. 1Bi**). The poor dispersion in the amide proton dimension of the HSQC (**Fig. 1B**) was a telltale sign that the CARD is (largely) unfolded. Additional evidence is obtained from 1D steady state ^15^N{^1^H} Nuclear Overhauser Effect (NOE) experiments that are sensitive to rapid picosecond-nanosecond (ps-ns) timescale dynamics (46). Average ^15^N{^1^H} NOE values based on 1D spectra recorded on CARD samples in buffer and in the presence of denaturing amounts of urea (6 M), -5°C, were identical, 0.52, supporting the notion of a largely unstructured CARD ensemble in buffer (*SI Appendix*, **Fig. S2A**).

For a well-structured protein, decreases in temperature often lead to a deterioration of NMR spectral quality as the overall tumbling of the molecule becomes slower as the temperature is lowered. Spectra of the CARD domain, however, show the opposite effect. This suggests that at 25°C, where very poor-quality amide correlation spectra are recorded, there is pervasive conformational heterogeneity on the microsecond-millisecond (μs-ms) timescale that leads to the deterioration in spectral quality. Conformational exchange was established over the complete temperature range examined, from 25°C to -5°C by recording a series of 1D ^15^N-based Carr-Purcell-Meiboom-Gill (CPMG) experiments (47). In **Figure 1Bii** a pair of 1D CPMG spectra recorded at -5°C is illustrated, showing an increase in peak intensities as a function of pulsing rate, ϖ_CPMG_, due to refocusing the effects of exchange. Additional experiments, recorded as pseudo 3D datasets at a pair of static magnetic fields (18.8 T and 11.7 T, -5°C) enabled a more quantitative, residue-specific analysis using a two-site model for chemical exchange where interconversion occurs between a minor (invisible) conformer and a major state, the latter giving rise to crosspeaks in spectra. Extracted values for the population of the minor state, *p_B_*, and the rate of exchange between the pair of states, *k_ex_*, varied between 1-5% and 200-700 s^-1^, respectively, (*SI Appendix*, **Fig. S2B**). Notably, the data could not be fit globally (i.e., all residues simultaneously), suggesting that the conformational heterogeneity derives from a set of non-cooperative interactions between different regions of the protein. The ^15^N-edited 1D spectra in **Figure 1Bi**, which show improved spectral quality with decreasing temperature, are consistent with decreases in *p_B_* or *k_ex_* or both from 25°C to -5°C. Unsurprisingly, motions in the μs-ms window were significantly reduced upon addition of 6 M urea, as the CPMG dispersion profiles became flat (*SI Appendix*, **Fig. 2B**).

**Figure 2.**
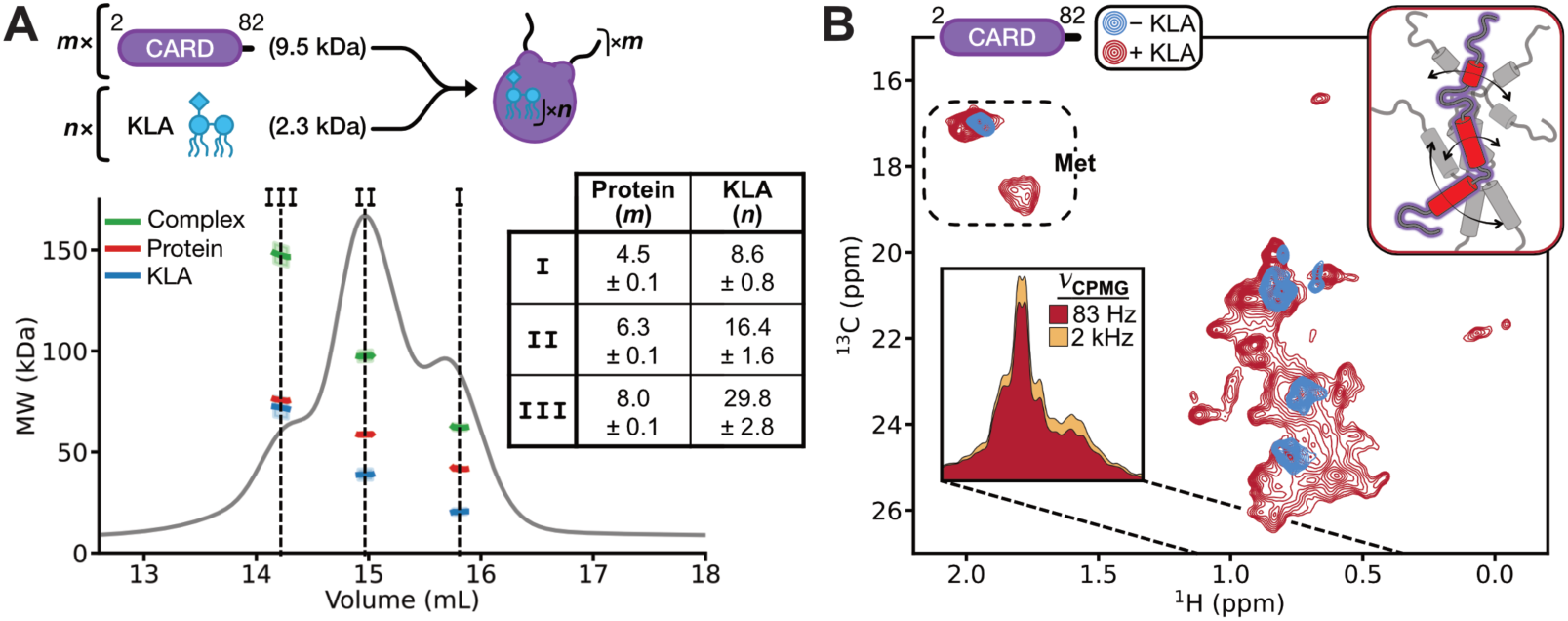
(A) SEC-UV-RI-LS data of Casp11 CARD in the presence of 3-fold molar excess KLA. A schematic for the stoichiometry of the non-canonical inflammasome scaffold under our *in-vitro* conditions is illustrated (top), with each complex consisting of *m* Casp11 protomers and *n* KLA molecules. A representative replicate of the SEC-MALS experiment is shown (bottom). The UV-trace (grey; y-axis not shown) indicates that three unique complexes can be separated (annotated as I, II, and III). Dotted black lines indicate positions where molecular weights were quantified to minimize contributions from adjacent peaks due to limited resolution. Using the ‘three-detector’ method (see *SI Appendix Text*) (53), the molecular weight of the complex (green), Casp11 CARD (red) and KLA (blue) can be elucidated. Errors from the uncertainty in the refractive index increment value of KLA are illustrated as lighter-shade bars for the complex and KLA values. Casp11 CARD and KLA stoichiometries are tabulated (right), with errors calculated using two experimental replicates. **(B)** 2D ^1^H-^13^C ddHMQC spectrum of U-^2^H, MLV-labelled Casp11 CARD prepared without (blue contours) or with 3-fold molar excess KLA (red contours). 1D ^13^C-edited MQ CPMG spectra recorded at either low (83 Hz; red-filled) or high ϖ_CPMG_ (2 kHz; yellow-filled) values are illustrated in the bottom-left inset. Data was recorded at 18.8 T (800 MHz, ^1^H frequency), 25°C. In the presence of KLA, the Casp11 CARD adopts a ‘molten globule’ state (59), with a cartoon depicting stable secondary structure (see (40) and *SI Appendix*, **Fig. S4** for CD data) and dynamic tertiary structure illustrated in the top-right inset.

^1^H-^13^C HSQC spectra of the isolated U-^13^C,^15^N CARD were also recorded (**Fig. 1C**), mirroring the peak intensity *vs.* temperature dependence observed for the amide data. As with the ^1^H-^15^N HSQC, the corresponding ^13^C spectrum was poorly dispersed (**Fig. 1C** focusing on the methyl correlations, right), again consistent with an unfolded polypeptide chain.

Having identified conditions where reasonable quality spectra could be recorded (−5°C), we sought to obtain assignments for the isolated CARD so that residue-specific structural dynamic insights could be obtained. Here we used a two-stage process. First, analysis of data from a suite of triple-resonance experiments (48, 49) resulted in assignments for approximately 90% of the residues (87% (68/78) NH, 91% (74/81) CO, 94% (76/81) CA and 86% (67/78) CB), limited by spectral quality due to conformational heterogeneity even at -5°C. Second, a sample of U-^13^C,^15^N CARD was prepared in 6M urea, -5°C and pH 6, and a complete set of assignments obtained as the resonances were no longer broadened (*SI Appendix*, **Fig. S3A**). Using a series of HNCO spectra (50), recorded on samples with progressively less urea, assignments were then transferred from denaturing to native conditions (*SI Appendix*, **Fig. S3B**), so that near complete assignments could be obtained (*SI Appendix*, **Fig. S3C,D**; 99% (77/78) NH, 99% (80/81) CO, 100% (81/81) CA and 90% (70/78) CB).

With assignments of the isolated Casp11 CARD available, the chemical shifts were then analyzed using the CheSPI program (51), to obtain residue specific populations of secondary structural elements. Notably, residues E24-F45 and A60-K63, from which the lowest intensity crosspeaks in ^1^H-^15^N spectra are derived (**Fig. 1D**; top), are predicted to have helix + turn populations exceeding 40% (**Fig. 1D**; bottom). The reduced peak intensities in this region are likely attributed to exchange between these conformations and unfolded populations (on a variety of timescales, including those outside of the CPMG window), leading to significant peak broadening.

The fraction of helix and turn in the CARD can also be established using CD spectroscopy, where α-helical and turn populations of 25% and 15%, respectively (*SI Appendix*, **Fig. S4**, blue-trace), are quantified, consistent with our NMR results. The α-helical content obtained is in agreement with a recent study using CD where the percent α-helix was estimated to be 16% (40). Taken together, our results establish that the Casp11 CARD is a highly dynamic motif, exchanging between unfolded and partially helical/turn elements of secondary structure (depicted by the cartoon illustration in **Figure 1D**), with a large region of the protein interconverting between conformers on the μs-ms timescale.

### The non-canonical inflammasome is highly heterogeneous

Prior to establishing whether the conformational heterogeneity of CARD in buffer persists when bound with LPS, we first prepared CARD : LPS complexes and determined the binding stoichiometry under *in-vitro* conditions favoring LPS micelle disaggregation (32). As described below (see ‘*Changes in KLA concentration affect the size distribution of non-canonical inflammasomes*’), this resulted in the formation of particle sizes on the order of 100 Å – 200 Å in diameter. We chose to incubate the Casp11 CARD with 3-fold molar excess of Kdo2-Lipid A (KLA), a truncated form of LPS that lacks the O-antigen and most of the core sugars (31, 52). KLA is significantly less heterogeneous than LPS and thus more favorable for biophysical studies, and previous reports indicate similar levels of activation of Casp11 with both KLA and LPS (29). Sample stoichiometry was established by the three detector SEC-UV-RI-LS method (53), whereby it was possible to extract both the total molecular mass of the complex, as well as the masses of the individual components within each complex. As the monomeric molecular masses of each of the components are known, it becomes possible to determine the stoichiometries of lipid and protein. Surprisingly, the resulting profile showed the presence of three different species, referred to as complexes I, II, and III, with the values of *m* and *n* for each of the three *m* CARD: *n* KLA complexes listed in the insert to **Figure 2A**. Notably, both *m* and *n* increase as the size of the complex grows, with the ratio of KLA to CARD molecules increasing as the overall molecular weight gets larger.

Attempts to perform similar experiments using inactive (C254A), full-length (FL) Casp11 were unsuccessful due to formation of minor amounts of soluble aggregates complicating analysis, and difficulties with resolving peaks II and III to enable quantitative determination of masses, as these peaks are less well separated than for the CARD : KLA complexes. This problem is exacerbated by a ‘broadening effect’ in three detector-based experiments where the peak resolution deteriorates with the transfer of solution to successive detectors. To ensure that similar results are obtained when a globular domain is attached to the C-terminus of CARD, as is the case for the PD of FL Casp11, we repeated these experiments with CARD-linker constructs fused to either Trp-Cage (30 amino acids), GB1 (64 amino acids), or SUMO (115 amino acids). The number of protein molecules associated with each of complexes I, II, and III was found to be independent of whether a globular domain was present or which of the three domains listed above was used, although interestingly there were some differences in the number of KLA molecules (*SI Appendix*, **Fig. S5A-D**). It should be mentioned that the presence of stoichiometric heterogeneity within each complex cannot be ruled out so that the values reported here are averages.

Having established the composition of CARD : KLA complexes, we next recorded NMR spectra to assess whether the structure of CARD becomes stabilized in a lipid environment. Previous CD studies of CARD in the presence of KLA demonstrated a secondary structure transition upon interaction with LPS, resulting in a doubling of the amount of alpha helicity to ∼50% (40), and we replicated these findings here (*SI Appendix*, **Fig. S4**, red-trace). As the molecular weight of the CARD : KLA complex is considerably larger than CARD alone, we choose to work with a highly deuterated protein for NMR studies, with protonation restricted (i) to only one of the two pro-chiral methyl groups of Leu and Val, labelled as ^13^CH_3_ (54), and (ii) to Met residues with ^13^C labelling only at the methyl position (55) (referred to in what follows as U-^2^H, MLV-labelling). Interestingly, we observed that the U-^2^H, MLV-labelled Casp11 CARD preferentially formed complex I, so that a monodisperse sample could be studied. ^1^H-^13^C delayed-decoupled heteronuclear multiple quantum coherence (ddHMQC) spectra (56) were recorded for both the isolated CARD as well as the CARD : KLA complex over a temperature range extending from 25°C to -5°C, pH 7.4 (**Fig. 2B**, recorded at 25°C). Although the spectral dispersion in the ^1^H-^13^C datasets improved in the presence of KLA (compare red *vs*. blue peaks) there remained significant peak broadening over all temperatures examined, and a noticeable increase in numbers of peaks from what was expected. Peak duplication is particularly apparent for Met residues where at least four resonances arise from the two methionines in the Casp11 CARD construct that we have used. As with the isolated CARD, and apparent from the quality of the spectrum, μs-ms timescale conformational dynamics are pervasive in the complex. This was confirmed by recording 1D ^13^C edited ^13^C-^1^H multiple-quantum (MQ) CPMG spectra (**Fig. 2B**, bottom-left inset) (57), showing an increase in peak intensities across much of the spectrum as ϖ_CPMG_ rates become larger.

Taken together, our results establish that although the percentage of helical structure is roughly doubled when CARD is mixed with KLA, the domain remains conformationally heterogeneous, with sidechains exchanging between states on the μs-ms timescale. In this context the ‘structure’ can be likened to that of a molten globule (58, 59), as depicted in the top-right corner of **Figure 2B**. Consistent with these observations, attempts to obtain structural data using cryo-EM were not successful due to the heterogeneous and dynamic nature of the complex (*SI Appendix*, **Fig. S6**).

### Changes in KLA concentration affect the size distribution of non-canonical inflammasomes

Having established three distinct classes of particles from preparations of either the isolated CARD domain or CARD-TrpCage/GB1/SUMO generated with 3-fold molar excess KLA, we wondered if a similar level of heterogeneity would be observed for FL Casp11 in the non-canonical inflammasome and, if so, how this would vary with the addition of different amounts of KLA. A previous study based on SEC showed that incubation of the Casp4 CARD or inactive (C258A) FL Casp4 with increasing amounts of LPS resulted in particles that increased in size with the amount of lipid added (32). However, under the conditions of these experiments the non-canonical inflammasomes eluted as only a single, broad peak, so that information on the individual complexes was not available. To establish whether the increased size reflected changes in relative amounts of each complex (I, II, and III) that we had observed in our studies, we incubated the C254A FL Casp11 with 0- to 5-fold molar equivalents of KLA and quantified the particle distributions using SEC (**Fig. 3A**, top). Under our conditions individual profiles of the three complexes could be resolved by deconvolution so that the amount of each component could be quantified as a function of KLA. In the absence of KLA, C254A FL Casp11 eluted from the SEC column as a monomer with notable peak tailing likely due to non-specific interactions of the CARD with the column matrix. Addition of increasing molar equivalents of KLA led to (i) the formation of non-canonical inflammasome complexes of varying sizes, (ii) changes of the populations of complexes I-III, and (iii) a concomitant decrease in the amount of monomer peak (**Fig. 3A**, top). Notably, under conditions of 3-fold or more excess KLA what appear to be soluble aggregates also began to form (**Fig. 3A**, grey areas under curve). By fitting the resulting SEC traces to a series of Gaussians the fraction of free Casp11 and of each complex could be determined at each concentration of KLA (**Fig. 3A**, bottom), showing an ordered transformation in the sizes of complexes – from small to large.

**Figure 3.**
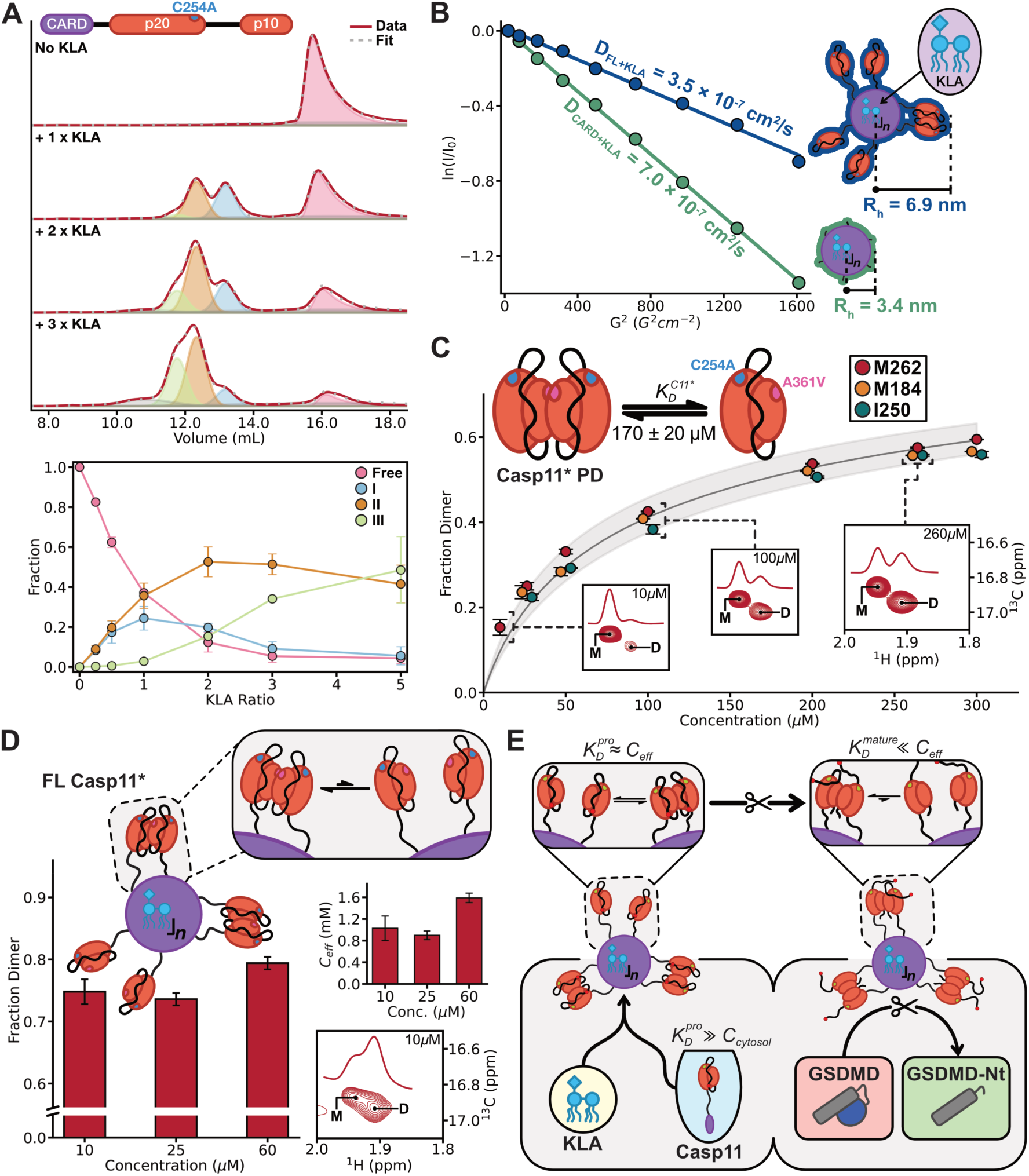
(A) (top) Representative size-exclusion profiles of C254A FL Casp11 (red lines) as a function of increasing amounts of KLA, written as molar equivalents of KLA to Casp11. Each peak is fit to either a Gaussian (I: blue, II: orange, III: green) or a skewed-Gaussian (Free: pink) lineshape. Two additional Gaussian functions, along with an additional skewed-Gaussian, are included in the fit to account for contaminants/aggregates (grey), with the summation of all fitted components represented by a dotted grey line. (bottom) The fraction of each fitted component (circles) is plotted as a function of KLA ratio, with errors calculated using two replicates; the line connecting experimental points is to guide the eye. **(B)** Diffusion constant measurements for Casp11 CARD (green) and C254A FL Casp11^PR^ (navy) with 3-fold molar excess KLA, acquired using methyl-based pulsed-field gradient NMR diffusion experiments (60). Extracted hydrodynamic radii are annotated on cartoon structures of complex II; the measured values are an average of all three complexes. **(C)** Casp11* PD fraction dimer as a function of concentration using relaxation corrected monomer and dimer peak-volumes measured in a 2D ^1^H-^13^C HSQC spectrum of a U-^1^H, IM-labelled sample. Spectral regions focussing on the M262 monomer-dimer pair are illustrated at different concentration in the insets, with 1D-traces of box-sums (^13^C ppm range: 16.75 – 17.05) shown. Fraction dimer values from each residue are displayed as filled circles, with error bars derived from standard deviations based on at least two measurements. The Met 184 and Ile 250 circles are shifted slightly to the left or right, respectively, for illustrative purposes. Curves from each residue were fit independently to a monomer-dimer dissociation model (𝑀 + 𝑀 ⇌ 𝐷), with the average fit (grey line) and 95%-confidence interval, based on the standard-deviation between the three fits (lighter-shading), displayed. The extracted 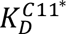 value is shown along with a cartoon depicting the monomer-dimer equilibrium. **(D)** Fraction dimer of the protease domain in the non-canonical inflammasome complex generated with FL Casp11* mixed with 3-fold molar excess KLA. Also shown is *C_eff_* of the protease domains (histogram in top right; see *SI Appendix Text* ‘*Estimating effective concentration of Casp11* in the non-canonical inflammasome*’). Analyses were based on the Met 262 monomer-dimer pair (spectrum of 10μM bulk Casp11* shown in bottom right). A schematic outlining the non-canonical inflammasome and protease domain dimerization is illustrated above. **(E)** Schematic of the Casp11 activation mechanism. In brief, pro-Casp11 is monomeric prior to interaction with KLA due to very weak dimerization affinity, 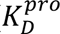. Formation of the non-canonical inflammasome allows pro-Casp11 dimers to form as 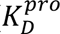 ≈ *C_eff_*, enabling proteolytic processing at Asp 285, and generating mature Casp11 whose dimerization affinity is strong, with 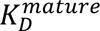 ≪ *C_eff_*. The dimeric, mature Casp11 can readily process GSDMD into its pore-forming, N-terminal domain (GSDMD-Nt), enabling an inflammatory response.

To provide an estimate of the effective size of the non-canonical inflammasome and establish whether its structure could be described in terms of a simple spherical model we used pulsed field gradient NMR diffusion measurements (60) of samples where either the Casp11 CARD (containing 3 additional Met residues on the C-terminus to improve signal intensity) or C254A FL Casp11^PR^ (PR = protease resistant, contains mutations to minimize degradation by trace amounts of contaminant proteases; see *SI Appendix Text*, ‘*Rationale for Casp11 mutations*’) was prepared with a 3-fold molar-excess of KLA (**Fig. 3B**). Although some of the mutations in C254A FL Casp11^PR^ were in the CARD, its ability to interact with LPS and form the non-canonical inflammasome remained intact, albeit with some changes in the overall distribution of complexes formed (*SI Appendix*, **Fig. S7**). As separate NMR resonances are not obtained for complexes I, II, and III, the experimental hydrodynamic radii (*R_h_*) values are averages over all particles. Diffusion coefficients of 3.5 ± 0.1 ×10^-7^ cm^2^/s and 7.0 ± 0.1 ×10^-7^ cm^2^/s were quantified for the complexes prepared with FL Casp11^PR^ and CARD, respectively, corresponding to *R_h_* values of 69 ± 2 Å and 34 ± 1 Å. To rationalize this difference, we constructed a simple model whereby the non-canonical inflammasome is formed from a central CARD : KLA scaffold (purple circle with blue KLA) with PDs (red ovals) projecting outwards (**Fig. 3B**, top cartoon). *R_h_* values of 12 Å and 24 Å are estimated for the ∼20-residue linker and the 271-residue PD components of the inflammasome using formulae relating *R_h_* to the number of residues in unfolded and folded proteins (61). Using these *R_h_* values along with the experimentally determined *R_h_* for the scaffold portion (34 Å), an effective *R_h_* for the non-canonical inflammasome of 70 Å is calculated, in excellent agreement with the measured value of 69 Å. Although a good match between experiment and the predicted *R_h_* is found using a simple spherical model description for an ‘average’ particle, this does not rule out the existence of other structural arrangements for the non-canonical inflammasome.

### Dimerization of the Casp11 PD is triggered by the formation of the non-canonical inflammasome

Initiator caspases, such as Casp9 involved in apoptosis, and inflammatory caspases, such as Casp4 and Casp11 that are critical for mounting an immune response in humans and mice, respectively, are monomeric in their inactive ‘pro-Casp’ states and become activated only upon dimerization of their respective PDs (34, 35, 62). Dimerization is also accompanied by auto-cleavage of an aspartic acid residue within the p20-p10 linker (Asp 285 for Casp11) to form the mature, active protease (34, 36). To establish the dimeric state of the PDs of FL Casp11 in the non-canonical inflammasome complex, and, hence, discern if the complex is in an active conformation even in the absence of substrates, the self-dimerization constant of the isolated PDs as well as their effective concentration (*C_eff_*) in the inflammasome particles must be known. The dimerization affinity of pro-Casp11 was determined by quantifying the relative intensities of well resolved crosspeaks for Ile 250 of the monomer and dimer forms in ^1^H-^13^C HSQC spectra as a function of protein concentration, taking into account differences in transverse relaxation rates of spins in the monomer and dimer (see *SI Appendix Text*, ‘*Correcting for differences in monomer-dimer peak transverse relaxation rates*’, ‘*Determination of K_D_ for Casp11* PD and catalytically active D277A, D285A Casp11 PD*’, & *SI Appendix*, **Fig. S8A**). Here we used a U-^1^H, Ile-labelled sample of catalytically active (Cys 254) D277A D285A Casp11 PD (residues 92-373). The Asp to Ala substitutions ensure that cleavage of the linker connecting p20-p10 cannot occur so that the protein is always in the pro-Casp form. The fraction dimer for all protein concentrations in the titration could be well fit to a monomer-dimer dissociation model, with a dissociation constant, 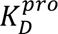, of 650 ± 50 µM.

In addition to measuring 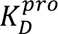 we also wanted to quantify the dimerization propensity of the mature form of the isolated PD where Asp 285 is cleaved, 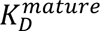. Because we suspected that 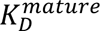 would be sub-μM, and therefore below the threshold for accurate quantification by NMR, we prepared an unlabelled sample of wild-type Casp11 PD (residues 92-373) and performed Analytical Ultracentrifugation Sedimentation Velocity (SV- AUC) experiments at a series of PD concentrations ranging from 0.1 to 19 μM. The data could be subsequently fit to a monomer-dimer dissociation model with a best fit 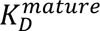 value of 0.66 μM, and a 68% confidence interval of 0. 26 µM ≤ 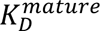 ≤ 8.65 µM (*SI Appendix*, **Fig. S8B**). Auto-cleavage of the Casp11 p20-p10 linker, thus, leads to a significant increase in the PD dimerization propensity.

To estimate *C_eff_*, we utilized a C254A FL Casp11 construct (catalytically dead, linker intact) that included several mutations to resist contaminant protease cleavage that was limiting otherwise (See *SI Appendix Text*, *‘Rationale for mutations’*). Additionally, an A361V mutation was added to increase the dimerization affinity and hence the intensities of dimer peaks in NMR studies of the non-canonical inflammasome. This is an important practical consideration as the large molecular weights of the complexes (ranging from ∼200 kDa to ∼450 kDa), and the low concentrations of particles used to ensure only intra-inflammasome PD interactions (bulk concentration of Casp11: 10 μM - 60 μM, concentration of particles ranging from ∼2 μM – 15 μM), challenged spectral sensitivity. In what follows this variant is referred to as Casp11*. As discussed previously (5), and further in the *SI Appendix Text*, ‘*Estimating effective concentration of Casp11* in the non-canonical inflammasome*’, *C_eff_* can be established once *K_D_* for the isolated Casp11* PDs (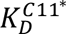) is known. 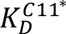 was obtained from a series of ^1^H-^13^C HSQC spectra of U-^1^H, IM-labelled Casp11* PD recorded at concentrations ranging from 10 to 300 μM. Using three well resolved monomer-dimer peak pairs (M184, I250, M262) assigned via a combination of mutagenesis (63) and magnetization-exchange experiments (63, 64) (*SI Appendix*, **Fig. S9**), relaxation-corrected peak volumes were used to calculate residue-specific fraction dimer values at each concentration that were subsequently fit to a monomer-dimer dissociation model, yielding 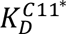 = 170 ± 20 µM (**Fig. 3C**).

To measure *C_eff_* we produced non-canonical inflammasomes comprised of U-^1^H, IM-labelled FL Casp11* with 3-fold molar excess of KLA. Three samples with bulk concentrations of FL Casp11* of 10 μM, 25 μM, and 60 μM were generated, and the fraction dimer was measured in a similar fashion as for the isolated Casp11* PD. The fraction dimer 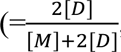, where [D] and [M] are the concentrations of monomer and dimer, respectively) was found to be largely invariant to the bulk concentration of Casp11*, and hence to the concentration of particles, as would be expected for intra-particle PD interactions, with an average value of 76 ± 3% across the concentration range examined (**Fig. 3D**, left). Calculated *C_eff_* values at each concentration of Casp11* are shown in **Figure 3D** (top right); an average value of 1170 ± 370 µM is obtained, which includes contributions from particles in complexes I, II, and III. Thus, the Casp11* PDs are largely dimeric in the context of the inflammasome, as illustrated by the cartoon in **Fig. 3D** (top). Importantly, when considered in the context of pro-Casp11 (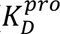 = 650 ± 50 µM), the dimeric form (as defined by fraction dimer) is also prevalent. Note that *C_eff_* could not be rigorously estimated using the mature form of Casp11 (linker cleaved), instead of Casp11*, as the single digit 𝜇M value for 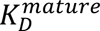 (∼1 μM) would ensure complete PD dimerization on the inflammasome for which only a lower estimate for *C_eff_* would then be obtained. **Figure 3E** shows a schematic of the Casp11 activation process based on the results presented here, leading to direct cleavage of GSDMD and downstream processing of pro-IL-18 and pro-IL-1β for signal propagation.

## Discussion

The non-canonical inflammasome is an essential component of the innate immune system, facilitating the detection of intracellular gram-negative bacteria and initiating an immune response against them (*SI Appendix*, **Fig. S1**). Although its benefits for the survival of the host are clear, dysregulation of the non-canonical inflammasome may lead to severe inflammatory diseases (25). Elucidating the structural features of this molecular machine including its mechanism of action, as a first step towards the development of therapeutics for modulating function in the case of severe inflammation are, therefore, important. Yet, even though the non-canonical inflammasome was discovered over a decade ago, and consists of only two major components, LPS and either Casp4 or Casp5 in humans or Casp11 in mice (27–29), detailed structural studies have not been forthcoming. Here we have used solution NMR spectroscopy, in concert with other biophysical techniques, to provide insights into the structural dynamics of these inflammasomes, their LPS/protein stoichiometry, and their mechanism of action.

Our NMR study establishes that while the Casp11 CARD is largely unstructured in solution, short regions of helical secondary structure are formed transiently, leading to pervasive dynamics on the μs-ms timescale (**Fig. 1**). Notably, the regions of transient structure established here align well with those identified in both Casp4 and Casp11 by HDX-MS, showing reduced deuterium uptake in the non-canonical inflammasome relative to the isolated CARD (40, 42), and suggesting that the transient helices become more prevalent in the context of the inflammasome complex. The NMR results highlighting a predominantly unstructured domain are consistent with CD studies by both our group and others (40), yet inconsistent with a previously published X-ray structure of Casp11 CARD in which a four-helix bundle structure is observed (37). It is likely that the maltose-binding protein (MBP) tag on the N-terminus of CARD, that was required for crystallization, led to stabilization of the non-physiological CARD structure through MBP-CARD contacts, as were noted in the original publication. Notably, the helix bundle CARD structure observed in the crystal was not formed upon addition of LPS, and although our study shows that helical content of the CARD increases to approximately 60%, it remains highly dynamic in the complex, with sidechains adopting multiple conformations.

Under conditions leading to LPS micelle disaggregation (32) we have established the CARD : KLA stoichiometry in the non-canonical inflammasome (**Fig. 2**). In a series of SEC-UV-RI-LS experiments we find that the resulting non-canonical inflammasome is composed of three unique complexes, containing either four, six or eight Casp11 CARDs, each with increasing stoichiometric amounts of KLA. While the numbers of CARD and lipid components obtained here represent the first quantitative measures in non-canonical inflammasomes, a recent study utilizing a combination of native mass spectrometry and mass photometry had identified the presence of distinct non-canonical inflammasome complexes and established each of their molecular masses (40). Notably, these masses could be validated by using the KLA : CARD stoichiometries determined for each complex in our study.

The 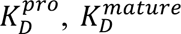, and *C_eff_* values that we have measured (**Fig. 3**) provide important insights into the mechanism of activation of the non-canonical inflammasome, as summarized in **Figure 3E**. Casp4 is constitutively expressed in humans, and it is estimated that it exists at a concentration of 10 - 60 nM in the cytosol (65, 66). Casp11 is not constitutively expressed and prior to induction via pro-inflammatory stimuli it cannot be detected in the cytosol (67). Although an accurate estimate of the Casp11 cytosolic concentration after induction has not been reported, it must be significantly less than the concentration of actin (20-100 μM), the most abundant family of proteins in eukaryotic cells (68, 69). As 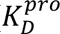 = 650 ± 50 µM >> [actin] >> [Casp11], Casp11 dimerization, and hence activation, can realistically only occur via formation of the non-canonical inflammasome. The results of this study, thus, strongly refute a recently proposed mechanism whereby Casp11 PD activation occurs prior to non-canonical inflammasome formation (70). Scaffolds, such as the non-canonical inflammasome, concentrate caspase PDs on their surface. Using non-canonical inflammasome particles prepared with FL Casp11* and 3-fold molar excess KLA, we quantified a PD effective concentration, *C_eff_*, of 1170 ± 370 µM, that is on the order of 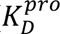. Thus, greater than one half of the Casp11 PDs would initially be dimerized in complexes, and, hence, activated through self-cleavage at residue Asp 285 in the p20-p10 linker. Note that activation through cleavage increases the dimerization propensity (0. 26 µM ≤ 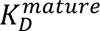 ≤ 8.65 µM) so that the PDs remain dimeric once cleavage at Asp 285 occurs.

The observation that the mature Casp11 PD exists almost exclusively as dimers 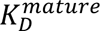 ≪ *C_eff_*) in the absence of substrate in the non-canonical inflammasome complex (**Fig. 3**) is distinct from what appears to occur on the surface of the apoptosome upon binding of its cognate caspase, Casp9 (5). There, dimerization requires substrate binding as the dimer dissociation constant for FL Casp9 is on the order of 15 mM, over an order of magnitude above the effective concentration of the PDs on the surface of the apoptosome (∼500 μM). Thus, even though the apoptosome concentrates Casp9 by over 7000-fold from its level in the cytosol, the fraction of Casp9 PD dimers would not exceed more than ∼5%. Why do systems which share such a fundamentally similar function exhibit this difference (**Fig. 4**)? One hypothesis is that the discrepancy may, in part, reflect differences in the cascades that the apoptosome and the inflammasome initiate. The sole purpose of apoptosis is to induce cell death without harming neighbouring cells. In contrast, the main function of pyroptosis is not to necessarily kill the cell, but instead to trigger an inflammatory response in neighbouring cells. This could necessitate a rapid release of pro-inflammatory cytokines that would be efficiently achieved through a fully ‘primed’ complex, as opposed to the case for the apoptosome where an extra layer of regulation is provided by requiring substrate binding for activation.

**Figure 4.**
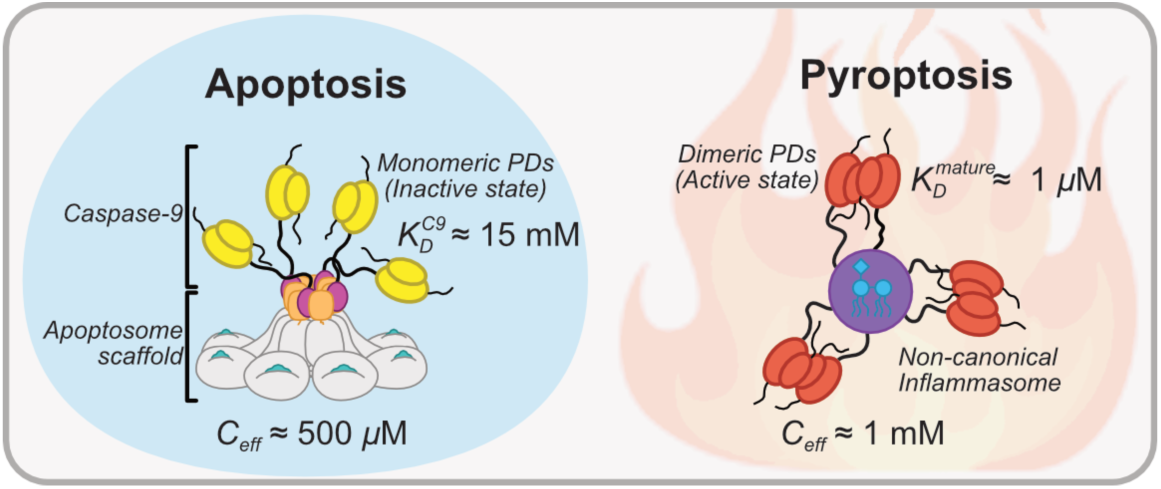
Comparison of protease domain dimerization equilibria in the apoptosome and the non-canonical inflammasome. While *C_eff_* ≥ 500μM for both apoptosome (5) and non-canonical inflammasomes, only the Casp11 PDs readily form the active dimeric state in the absence of substrate due to a significantly lower 𝐾*_D_* value.

In summary, our study paints a picture of the non-canonical inflammasome as a highly dynamic and heterogenous structural ensemble that facilitates rapid activation of Casp11 to promote the release of cytokines (**Fig. 4**). This is achieved by creating an effective concentration of Casp11 molecules on the inflammasome surface that favors PD dimerization and, hence, activation. Our work further highlights the dynamic nature of molecular machines, emphasizing the important role that solution NMR spectroscopy can play in understanding how these systems function (71).

## Materials and Methods

### Cloning, expression, purification, and sample preparation

All constructs utilized in this study were codon-optimized (see *SI Appendix*, Table S1) and sub-cloned into pET vectors for overexpression in an *E. coli* host. Proteins were purified to homogeneity using routine purification methods. Detailed descriptions of all DNA constructs, expression, purification, and sample preparation are provided in the *SI Appendix Text*, ‘*Materials and Methods*’.

### Data acquisition and analysis

NMR experiments were recorded at either 23.5 Tesla (1 GHz ^1^H frequency Bruker Avance NEO), 18.8 Tesla (800 MHz Bruker Avance III HD), 14.1 Tesla (600 MHz Bruker Avance III HD), or 11.7 Tesla (500 MHz Bruker Avance III HD) using spectrometers equipped with either cryogenically cooled x, y, z axis pulsed-field gradient triple-resonance probes (23.5 T, 18.8 T, 14.1 T) or a liquid nitrogen cooled z axis pulse-field gradient triple resonance probe (11.7 T). SEC-UV-RI-LS and SEC-UV measurements were acquired using a Superdex 200 Increase 10/300 GL column coupled to an Agilent 1260 Infinity II HPLC with options for in-line UV-vis (set to 280 nm), RI (Wyatt Optilab T-rEX) and LS (Wyatt DAWN) detectors and an autosampler. CD measurements were performed using a Jasco J-1500 spectrophotometer with 1 mm quartz cuvettes. Cryo-EM data were acquired using a Thermo Fischer Glacios electron microscope operating at 200 kV and equipped with a Falcon 4i direct detector device camera. SV-AUC measurements were performed on a Beckman Coulter ProteomeLab XL-I ultracentrifuge. For specific sample conditions see *SI Appendix*, Table S2, and for a detailed description of all experimental setups and data analysis, see *SI Appendix Text*, *‘Materials and Methods’*.

## Author Contributions

Project conception and design of research: A.I.M.S. and L.E.K.; Sample production: A.I.M.S.; NMR experiments and data analysis: A.I.M.S., J.M.A., J.P.B. and L.E.K.; SEC-UV-RI-LS, SEC, and CD experiments and data analysis: A.I.M.S.; AUC experiments and data analysis: H.Z. and P.S.; Cryo-EM experiments and data analysis: H.W. and J.L.R. Paper writing: A.I.M.S. and L.E.K. with input from all authors.

## Data Availability

All data are included in the article and/or *SI Appendix*.

## Acknowledgements

This work was supported through grants from the Canadian Institutes of Health Research (CIHR; FND-503573 to L.E.K.), the Natural Sciences and Engineering Council of Canada (NSERC; 024-03872 to L.E.K.), and Intramural Research Programs of NIBIB (ZIA EB000099-02 to P.S.) at the National Institutes of Health (NIH). A.I.M.S. is grateful to NSERC for a Canada Graduate Scholarship – Doctoral, J.P.B. acknowledges funding from a CIHR Post-Doctoral Fellowship, and H.W. holds a Mary H. Beatty Fellowship. The contributions of the NIH affiliated authors (H.Z. and P.S.) are considered Works of the United States Government. The findings and conclusions presented in this paper are those of the authors and do not necessarily reflect the views of the NIH or the U.S. Department of Health and Human Services. The authors thank the staff of the Structural and Biophysical Core Facility of the Hospital for Sick Children for assistance in SEC-UV-RI-LS data collection, and Prof. T. Reid Alderson (Helmholtz Munich), Prof. M. Stephen Trent (University of Georgia), and Prof. Russel Bishop (McMaster University) for helpful discussions.

## References

1. J. Yuan, D. Ofengeim, A guide to cell death pathways. Nat. Rev. Mol. Cell Biol. 25, 379–395 (2024).

2. Y. Li, et al., Mechanistic insights into caspase-9 activation by the structure of the apoptosome holoenzyme. Proc. Natl. Acad. Sci. 114, 1542–1547 (2017).

3. T. C. Cheng, C. Hong, I. V. Akey, S. Yuan, C. W. Akey, A near atomic structure of the active human apoptosome. eLife 5, e17755 (2016).

4. M. Zhou, et al., Atomic structure of the apoptosome: mechanism of cytochrome c - and dATP-mediated activation of Apaf-1. Genes Dev. 29, 2349–2361 (2015).

5. A. I. M. Sever, et al., Activation of caspase-9 on the apoptosome as studied by methyl-TROSY NMR. Proc. Natl. Acad. Sci. 120, e2310944120 (2023).

6. R. Chen, et al., Pattern recognition receptors: function, regulation and therapeutic potential. Signal Transduct. Target. Ther. 10, 216 (2025).

7. P. Broz, V. M. Dixit, Inflammasomes: mechanism of assembly, regulation and signalling. Nat. Rev. Immunol. 16, 407–420 (2016).

8. N. A. Thornberry, et al., A novel heterodimeric cysteine protease is required for interleukin-1β processing in monocytes. Nature 356, 768–774 (1992).

9. T. Ghayur, et al., Caspase-1 processes IFN-γ-inducing factor and regulates LPS-induced IFN- γ production. Nature 386, 619–623 (1997).

10. F. Martinon, K. Burns, J. Tschopp, The Inflammasome. Mol. Cell 10, 417–426 (2002).

11. J. Shi, et al., Cleavage of GSDMD by inflammatory caspases determines pyroptotic cell death. Nature 526, 660–665 (2015).

12. G. Du, et al., ROS-dependent S-palmitoylation activates cleaved and intact gasdermin D. Nature 630, 437–446 (2024).

13. L. Sborgi, et al., GSDMD membrane pore formation constitutes the mechanism of pyroptotic cell death. EMBO J. 35, 1766–1778 (2016).

14. P. Devant, J. C. Kagan, Molecular mechanisms of gasdermin D pore-forming activity. Nat. Immunol. 24, 1064–1075 (2023).

15. C. L. Evavold, et al., The Pore-Forming Protein Gasdermin D Regulates Interleukin-1 Secretion from Living Macrophages. Immunity 48, 35–44.e6 (2018).

16. R. Heilig, et al., The Gasdermin-D pore acts as a conduit for IL-1β secretion in mice. Eur. J. Immunol. 48, 584–592 (2018).

17. S. Xia, et al., Gasdermin D pore structure reveals preferential release of mature interleukin-1. Nature 593, 607–611 (2021).

18. T. Jin, et al., Structures of the HIN Domain:DNA Complexes Reveal Ligand Binding and Activation Mechanisms of the AIM2 Inflammasome and IFI16 Receptor. Immunity 36, 561–571 (2012).

19. L. Sborgi, et al., Structure and assembly of the mouse ASC inflammasome by combined NMR spectroscopy and cryo-electron microscopy. Proc. Natl. Acad. Sci. 112, 13237–13242 (2015).

20. A. Lu, et al., Unified Polymerization Mechanism for the Assembly of ASC-Dependent Inflammasomes. Cell 156, 1193–1206 (2014).

21. L. Xiao, V. G. Magupalli, H. Wu, Cryo-EM structures of the active NLRP3 inflammasome disc. Nature 613, 595–600 (2023).

22. R. E. Matico, et al., Structural basis of the human NAIP/NLRC4 inflammasome assembly and pathogen sensing. Nat. Struct. Mol. Biol. 31, 82–91 (2024).

23. L. R. Hollingsworth, et al., Mechanism of filament formation in UPA-promoted CARD8 and NLRP1 inflammasomes. Nat. Commun. 12, 189 (2021).

24. K. P. Downs, H. Nguyen, A. Dorfleutner, C. Stehlik, An overview of the non-canonical inflammasome. Mol. Aspects Med. 76, 100924 (2020).

25. E. Elkayam, F. G. Gervais, H. Wu, M. A. Crackower, J. Lieberman, New insights into the noncanonical inflammasome point to caspase-4 as a druggable target. Nat. Rev. Immunol. (2025). 10.1038/s41577-025-01142-9.

26. N. Kayagaki, et al., Non-canonical inflammasome activation targets caspase-11. Nature 479, 117–121 (2011).

27. N. Kayagaki, et al., Noncanonical Inflammasome Activation by Intracellular LPS Independent of TLR4. Science 341, 1246–1249 (2013).

28. J. A. Hagar, D. A. Powell, Y. Aachoui, R. K. Ernst, E. A. Miao, Cytoplasmic LPS Activates Caspase-11: Implications in TLR4-Independent Endotoxic Shock. Science 341, 1250–1253 (2013).

29. J. Shi, et al., Inflammatory caspases are innate immune receptors for intracellular LPS. Nature 514, 187–192 (2014).

30. N. Kayagaki, et al., Caspase-11 cleaves gasdermin D for non-canonical inflammasome signalling. Nature 526, 666–671 (2015).

31. C. R. H. Raetz, C. Whitfield, Lipopolysaccharide Endotoxins. Annu. Rev. Biochem. 71, 635–700 (2002).

32. J. An, et al., Caspase-4 disaggregates lipopolysaccharide micelles via LPS-CARD interaction. Sci. Rep. 9, 826 (2019).

33. S. Wang, et al., Identification and Characterization of Ich-3, a Member of the Interleukin-1β Converting Enzyme (ICE)/Ced-3 Family and an Upstream Regulator of ICE. J. Biol. Chem. 271, 20580–20587 (1996).

34. C. Ross, A. H. Chan, J. Von Pein, D. Boucher, K. Schroder, Dimerization and auto-processing induce caspase-11 protease activation within the non-canonical inflammasome. Life Sci. Alliance 1, e201800237 (2018).

35. A. H. Chan, et al., Caspase-4 dimerisation and D289 auto-processing elicit an interleukin-1β-converting enzyme. Life Sci. Alliance 6, e202301908 (2023).

36. B. L. Lee, et al., Caspase-11 auto-proteolysis is crucial for noncanonical inflammasome activation. J. Exp. Med. 215, 2279–2288 (2018).

37. M. Liu, et al., Crystal structure of caspase-11 CARD provides insights into caspase-11 activation. Cell Discov. 6, 70 (2020).

38. J. Jumper, et al., Highly accurate protein structure prediction with AlphaFold. Nature 596, 583–589 (2021).

39. M. Varadi, et al., AlphaFold Protein Structure Database: massively expanding the structural coverage of protein-sequence space with high-accuracy models. Nucleic Acids Res. 50, D439–D444 (2022).

40. C. Wang, et al., LPS-induced structural reorganization and polymerization drive noncanonical inflammasome activation. Sci. Adv. (2026).

41. M. A. Wacker, A. Teghanemt, J. P. Weiss, J. H. Barker, High-affinity caspase-4 binding to LPS presented as high molecular mass aggregates or in outer membrane vesicles. Innate Immun. 23, 336–344 (2017).

42. J. Began, et al., Caspase-4 binds to LPS membranes with positive curvature for non-canonical inflammasome activation. Immunity 59, 542–558.e8 (2026).

43. K. Wang, et al., Structural Mechanism for GSDMD Targeting by Autoprocessed Caspases in Pyroptosis. Cell 180, 941–955.e20 (2020).

44. T. R. Alderson, L. E. Kay, NMR spectroscopy captures the essential role of dynamics in regulating biomolecular function. Cell 184, 577–595 (2021).

45. H. J. Dyson, P. E. Wright, NMR illuminates intrinsic disorder. Curr. Opin. Struct. Biol. 70, 44–52 (2021).

46. L. E. Kay, D. A. Torchia, A. Bax, Backbone dynamics of proteins as studied by nitrogen-15 inverse detected heteronuclear NMR spectroscopy: application to staphylococcal nuclease. Biochemistry 28, 8972–8979 (1989).

47. D. F. Hansen, P. Vallurupalli, L. E. Kay, An Improved^15^ N Relaxation Dispersion Experiment for the Measurement of Millisecond Time-Scale Dynamics in Proteins. J. Phys. Chem. B 112, 5898–5904 (2008).

48. M. Sattler, C. Griesinger, J. Schleucher, Heteronuclear multidimensional NMR experiments for the structure determination of proteins in solution employing pulsed field gradients. Prog. Nucl. Magn. Reson. Spectrosc. 34, 93–158 (1999).

49. S. C. Panchal, N. S. Bhavesh, R. V. Hosur, Improved 3D triple resonance experiments, HNN and HN(C)N, for H^N^ and ^15^N sequential correlations in (^13^C, ^15^N) labeled proteins: Application to unfolded proteins. J. Biomol. NMR 20, 135–147 (2001).

50. M. Ikura, L. E. Kay, A. Bax, A novel approach for sequential assignment of proton, carbon-13, and nitrogen-15 spectra of larger proteins: heteronuclear triple-resonance three-dimensional NMR spectroscopy. Application to calmodulin. Biochemistry 29, 4659–4667 (1990).

51. J. T. Nielsen, F. A. A. Mulder, CheSPI: chemical shift secondary structure population inference. J. Biomol. NMR 75, 273–291 (2021).

52. C. R. H. Raetz, et al., Kdo2-Lipid A of Escherichia coli, a defined endotoxin that activates macrophages via TLR-4. J. Lipid Res. 47, 1097–1111 (2006).

53. D. J. Slotboom, R. H. Duurkens, K. Olieman, G. B. Erkens, Static light scattering to characterize membrane proteins in detergent solution. Methods 46, 73–82 (2008).

54. V. Tugarinov, L. E. Kay, An Isotope Labeling Strategy for Methyl TROSY Spectroscopy. J. Biomol. NMR 28, 165–172 (2004).

55. I. Gelis, et al., Structural Basis for Signal-Sequence Recognition by the Translocase Motor SecA as Determined by NMR. Cell 131, 756–769 (2007).

56. N. Bolik-Coulon, et al., Less is more: A simple methyl-TROSY based pulse scheme offers improved sensitivity in applications to high molecular weight complexes. J. Magn. Reson. 346, 107326 (2023).

57. D. M. Korzhnev, K. Kloiber, L. E. Kay, Multiple-Quantum Relaxation Dispersion NMR Spectroscopy Probing Millisecond Time-Scale Dynamics in Proteins: Theory and Application. J. Am. Chem. Soc. 126, 7320–7329 (2004).

58. D. Barrick, R. L. Baldwin, The molten globule intermediate of apomyoglobin and the process of protein folding. Protein Sci. 2, 869–876 (1993).

59. C. Redfield, Using nuclear magnetic resonance spectroscopy to study molten globule states of proteins. Methods 34, 121–132 (2004).

60. W.-Y. Choy, et al., Distribution of molecular size within an unfolded state ensemble using small-angle X-ray scattering and pulse field gradient NMR techniques. J. Mol. Biol. 316, 101–112 (2002).

61. D. K. Wilkins, et al., Hydrodynamic Radii of Native and Denatured Proteins Measured by Pulse Field Gradient NMR Techniques. Biochemistry 38, 16424–16431 (1999).

62. M. Renatus, H. R. Stennicke, F. L. Scott, R. C. Liddington, G. S. Salvesen, Dimer formation drives the activation of the cell death protease caspase 9. Proc. Natl. Acad. Sci. 98, 14250–14255 (2001).

63. R. Sprangers, L. E. Kay, Quantitative dynamics and binding studies of the 20S proteasome by NMR. Nature 445, 618–622 (2007).

64. T. L. Religa, R. Sprangers, L. E. Kay, Dynamic Regulation of Archaeal Proteasome Gate Opening As Studied by TROSY NMR. Science 328, 98–102 (2010).

65. M. Beck, et al., The quantitative proteome of a human cell line. Mol. Syst. Biol. 7, 549 (2011).

66. M. Y. Hein, et al., A Human Interactome in Three Quantitative Dimensions Organized by Stoichiometries and Abundances. Cell 163, 712–723 (2015).

67. R. Schauvliege, J. Vanrobaeys, P. Schotte, R. Beyaert, Caspase-11 gene expression in response to lipopolysaccharide and interferon-gamma requires nuclear factor-kappa B and signal transducer and activator of transcription (STAT) 1. J. Biol. Chem. 277, 41624–41630 (2002).

68. R. Dominguez, K. C. Holmes, Actin Structure and Function. Annu. Rev. Biophys. 40, 169–186 (2011).

69. T. D. Pollard, L. Blanchoin, R. D. Mullins, Molecular Mechanisms Controlling Actin Filament Dynamics in Nonmuscle Cells. Annu. Rev. Biophys. Biomol. Struct. 29, 545–576 (2000).

70. D. C. Akuma, et al., Catalytic activity and autoprocessing of murine caspase-11 mediate noncanonical inflammasome assembly in response to cytosolic LPS. eLife 13, e83725 (2024).

71. A. I. M. Sever, R. Ahmed, P. Rößler, L. E. Kay, Solution NMR goes big: Atomic resolution studies of protein components of molecular machines and phase-separated condensates. Curr. Opin. Struct. Biol. 90, 102976 (2025).

