## Supplementary material for "The dynamic and heterogeneous structure of the non-canonical inflammasome": SI-Appendix+SI-Figures

### Materials and Methods

#### *Plasmid constructs and cloning*

The constructs utilized in this study were codon-optimized for overexpression in an *E. coli* host. All inactive (C254A) full-length (FL) Casp11 constructs (residues 1-373, UniProt: P70343), including the protease resistant (PR) variant used for diffusion measurements of the non-canonical inflammasome (Casp11$^{PR}$), and Casp11* which includes the PR mutations, A361V, and a number of additional Met residue mutations (see '*Rationale for Casp11 mutations'* below), were synthesized and sub-cloned into pET-IDT vectors (IDT; www.idtdna.com). All FL Casp11 variants are referred to collectively as FL Casp11$^{All}$ in what follows. The Casp11 CARD-only (residues 1-82), and Casp11 CARD-TrpCage/GB1/SUMO constructs were similarly cloned into pET-IDT vectors. Additional mutations to any of the constructs were introduced via QuikChange site-directed mutagenesis (Agilent).

The FL Casp11$^{All}$ constructs were synthesized with an N-terminal His$_6$-TEV tag, while the Casp11 CARD-only constructs were synthesized with a C-terminal SNAC-His$_6$-TwinStrep tag (1, 2). Casp11 CARD-TrpCage/GB1/SUMO (herein referred to collectively as CARD-fusion) constructs were synthesized with a non-cleavable C-terminal His$_6$-tag. Both catalytically active (Cys 254) and inactive (C254A) protease domain-only (PD) Casp11 (residues 92-373) constructs were synthesized by IDT as gBlocks and were sub-cloned via Gibson Assembly (New England BioLabs Inc.) into a pET vector containing an N-terminal His$_6$-MBP-TEV tag. C254A Casp11 PD constructs made use of a pET vector containing mutations in the T7 promoter and translation initiation region as described previously (3). The DNA and corresponding amino acid sequences for the major constructs utilized in this study are outlined in *SI Appendix,* Table S1, along with extinction coefficients (predicted by ExPASY ProtParam, https://web.expasy.org/protparam/) used to estimate protein concentration.

***Casp11 expression and purification***

All samples were produced via overexpression in *E. coli* and were grown in M9 minimal media. To produce unlabelled protein, FL Casp11$^{All}$, CARD-only, and CARD-fusion constructs were transformed into an LPS-deficient strain of *E. coli*, *ClearColi* BL21(DE3) (https://clearcoli.com/), and grown in M9 minimal media supplemented with 10 g/L NaCl. Similarly, C254A Casp11 PD and catalytically active Casp11 PD constructs were transformed into *E. coli* BL21(DE3)pLysS and BL21-CodonPlus (DE3)-RIPL cells (Agilent), respectively, and grown in M9 minimal media with 0.5 g/L NaCl. In all cases, the cells were initially grown at 37°C until $OD_{600}$ ~0.8 was reached and then the temperature was lowered to 18°C. Protein expression was subsequently induced 1 hour later with either 0.2 mM (*ClearColi* BL21(DE3), BL21-CodonPlus (DE3)-RIPL) or 0.5mM (BL21(DE3)pLysS) isopropyl β-D-1-thiogalactopyranoside (IPTG). The cultures were harvested 16-20 hours after induction and the pellet was placed at -20°C until purification. For production of $^{15}$N-labelled and/or $^{13}$C-labelled samples, cells were grown in $H_2O$ M9 minimal media containing $^{15}NH_4Cl$ and/or uniform-$^{13}$C-glucose as the sole nitrogen and carbon sources, respectively. For U-$^{1}$H samples with Ile or Met $^{13}CH_3$-labelling, precursors (60 mg/L α-ketobutyric acid, methyl-$^{13}$C for Ileδ1-$^{13}CH_3$; 100 mg/L methyl-$^{13}CH_3$-methionine for Metε-$^{13}CH_3$) were added 1 hour prior to induction of protein overexpression. Finally, for production of U-$^{2}$H samples with Ile, Leu, Val, or Met $^{13}CH_3$-labelling (with only one of the pair of isopropyl methyls of Leu and Val randomly labelled as $^{13}CH_3$), cells were grown in $D_2O$ M9 minimal media containing D-glucose-1,2,3,4,5,6,6-$d_7$ as the sole carbon source, and precursors (60 mg/L α-ketobutyric acid, methyl-$^{13}$C{3,3-$D_2$} for Ileδ1-$^{13}CH_3$; 100 mg/L α-ketoisovaleric acid, 3-methyl-$^{13}$C{3,4,4,4-$D_4$} for Leuδ,Valγ-$^{13}CH_3$/$^{12}CD_3$; and 100 mg/L methyl-$^{13}CH_3$-methionine for Metε-$^{13}CH_3$) were added 1 hour prior to induction of protein overexpression.

All purification protocols outlined below are for 1L of culture volume. The purification of all Casp11 CARD-only constructs made use of an inclusion body (IB) protocol. Cells were resuspended in lysis buffer (50 mM Tris, 300 mM NaCl, 3 mM β-mercaptoethanol (BME), pH 8.0) + 0.1 mg/mL DNase I + 0.1 mg/mL lysozyme, followed by sonication and then centrifugation at 14,000×g for 1 hour at 4°C. The supernatant was discarded, and the IB pellet was dissolved in 20 mL of resolubilization buffer (6 M Guanidine Hydrochloride (GdnHCl), 100 mM Sodium Phosphate (NaPhos), 500 mM NaCl, 10 mM imidazole, 0.5% v/v Tween-20, 3 mM BME, pH 8.0). The sample was then clarified via centrifugation and passed through a 1.0 µm syringe filter. The clarified, resolubilized IBs were then loaded onto a 5mL HisTrap HP column (Cytiva) using a benchtop peristaltic pump and then washed with 3× column volumes (CV) of resolubilization buffer. The column was then washed further with 6×CV of IB wash buffer (6 M GdnHCl, 100 mM NaPhos, 500 mM NaCl, 12.5 mM imidazole, 3 mM BME, pH 8.0), and then eluted with 5×CV of IB elution buffer (6 M GdnHCl, 100 mM NaPhos, 500 mM NaCl, 200 mM imidazole, 3 mM BME, pH 8.0). Subsequently, the eluant was buffer-exchanged (>1000×) into SNAC-cleavage buffer (5.5 M GdnHCl, 25 mM HEPES, 150 mM NaCl, pH 8.2) + 100 mM Acetone Oxime + 1 mM $NiCl_2$ and the final volume adjusted so that the protein concentration was ~1 mg/mL (1). The sample was incubated at 37°C for 16 hours to allow for complete cleavage of the SNAC-$His_6$-TwinStrep tag. After incubation, 8 mM imidazole was added to the sample, and this was passed through a 5mL HisTrap HP column followed by 3×CV of SNAC-cleavage buffer + 8 mM imidazole to remove the cleaved purification tag. The column flow-through was then concentrated so that the protein concentration was ~0.8–1.2 mM. Reconstitution of this sample into native buffer could then be achieved via dropwise dilution of 0.5 mL into 5 mL of size-exclusion chromatography (SEC) buffer (see *SI Appendix,* Table S2 for sample specific buffer composition) with rapid stirring (11×

dilution). The resulting solution was loaded directly onto a HiLoad 16/600 Superdex 75 PG column (Cytiva) using an FPLC and fractions containing the protein were collected and pooled. If required, the sample was diluted to achieve the correct analysis buffer conditions (see *SI Appendix,* Table S2), and was then subsequently concentrated to the desired concentration (between 75–125 μM) using a stirred cell concentrator with a 3-kDa molecular weight cutoff (MWCO), 25 mm diameter PolyEtherSulfone (PES) membrane. Notably, this protocol can yield tens of milligrams of protein per liter of culture. To ensure that the CARD was folded correctly using this protocol 2D $^{1}$H-$^{15}$N HSQC spectra (4) were recorded and compared with spectra of CARD prepared using a published protocol (5) which lacked denaturant:

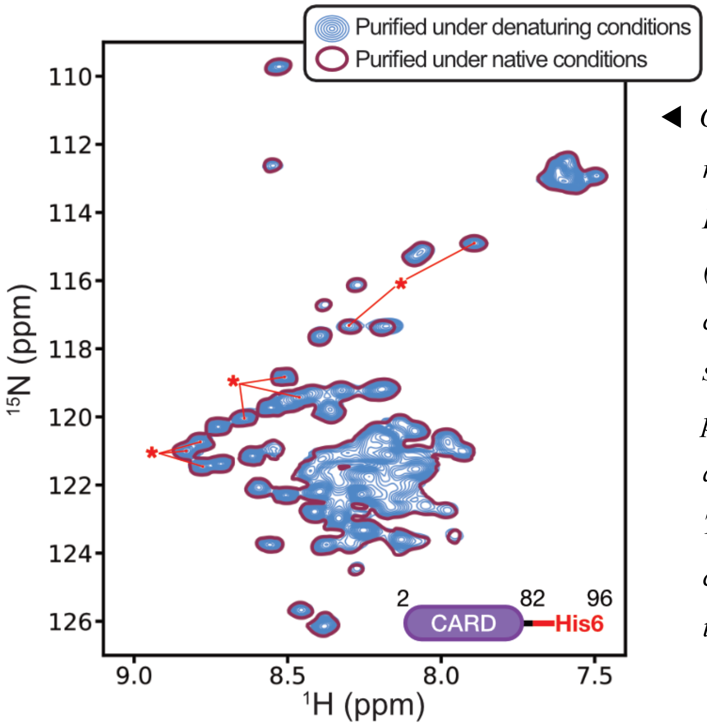


◀ *Overlay of 2D $^{1}$H-$^{15}$N HSQC spectra recorded on samples of Casp11 CARD-His6 purified using either 6 M GdnHCl (denaturing conditions, blue multi-contours) or under native conditions (red single-contours). Both samples were prepared in identical buffers and spectra acquired under identical conditions, 11.7 T (500 MHz) and -5 ℃. The '*' indicates crosspeaks which arise from the C-terminal linker+His6-tag.*

Purification of Casp11 CARD-fusion proteins was carried out with a simplified version of the CARD-only protocol with the following changes: 1) the cell pellet was resuspended directly in 20 mL of resolubilization buffer, skipping the IB wash step, and 2) the eluant from the nickel column was immediately concentrated to ~0.8–1.2 mM, and placed at -20°C until required, omitting the

SNAC-cleavage and second nickel column steps. Prior to reconstitution in non-denaturing conditions, 10 mM BME was added to the sample to ensure all cysteines were reduced. Notably, due to the absence of an N-terminal fusion-tag, the N-terminal methionines of the CARD-only and CARD-fusion constructs are removed via post-translational processing in the *E. coli* expression system.

Purification of FL Casp11$^{All}$ proteins was achieved with resuspension of protein pellets in 30mL of lysis buffer + 0.1 mg/mL DNase I + 0.1 mg/mL lysozyme + 20 mM imidazole + 1% Tween-20, followed by sonication and centrifugation. The supernatant was then filtered with a 1.0 µm syringe filter, and subsequently loaded onto a 1mL HisTrap HP column (Cytiva) using a benchtop peristaltic pump. The column was washed with 15×CV of lysis buffer + 40 mM imidazole + 1% Tween-20, and then 20×CV of lysis buffer + 40 mM imidazole + 200 mM NaCl (500 mM total). The column was then connected to an FPLC, and the protein was eluted with an imidazole gradient (50–300 mM). Fractions containing the protein were collected and 0.25 mg of hyperTEV60 protease (6) was added. The sample was dialyzed overnight into dialysis buffer (50 mM Tris, 400 mM NaCl, 3 mM BME, pH 8.0) at 4°C. Subsequently, the imidazole concentration in the sample was adjusted to 25 mM and it was passed again through a 1mL HisTrap HP column, followed by 10×CV of dialysis buffer containing 25 mM imidazole, in order to remove the purification tag and hyperTEV60. The column flow-through was then concentrated to 2–4 mL using a stirred cell concentrator with a 30-kDa MWCO, 25 mm diameter PES membrane. The concentrated solution was subsequently loaded directly onto a HiLoad 16/600 Superdex 75 PG column, equilibrated in SEC buffer (see *SI Appendix,* Table S2), and fractions containing the protein were collected and pooled. The sample was then diluted to obtain the correct analysis buffer conditions, and then concentrated to the desired concentration using a stirred cell concentrator.

Finally, C254A Casp11 PD and catalytically active Casp11 PD samples were purified by resuspension of cell pellets in lysis buffer + 0.1 mg/mL DNase I + 0.1 mg/mL lysozyme + 25 mM imidazole followed by sonication and centrifugation. The supernatant was filtered using a 1.0 µm syringe filter, and subsequently loaded onto a 5mL HisTrap HP column using a benchtop peristaltic pump. The column was then washed with 10×CV of lysis buffer + 200 mM NaCl (500 mM total) + 25 mM imidazole and eluted with 5×CV of lysis buffer + 500 mM imidazole. 0.5 mg hyperTEV60 protease (6) was added to the eluant, followed by dialysis overnight into dialysis buffer at 4°C. Subsequently, the imidazole concentration in the sample was adjusted to 20 mM and the sample was passed through a 5mL HisTrap HP column, followed by a 3×CV wash with dialysis buffer containing 20 mM imidazole. The column flow-through was then concentrated to 2–4 mL using a 10-kDa MWCO centrifugal spin concentrator, and then loaded onto a HiLoad 16/600 Superdex 75 PG column equilibrated with 50 mM Tris, 150 mM NaCl, 5 mM dithiothreitol (DTT), pH 8.0. Fractions containing the protein were collected and buffer-exchanged (>1000×) into the desired analysis buffer (see *SI Appendix,* Table S2) using a 10-kDa MWCO centrifugal spin concentrator at 4ºC.

***Preparation of Casp11 : KLA complexes***

FL Casp11$^{All}$, CARD-only, and CARD-fusion proteins were purified as described in the previous section. Kdo2-Lipid A (KLA) (7) was purchased from Avanti Research (https://www.avantiresearch.com/) in the form of a lyophilized powder (1mg, Cat# 699500) and resuspended in 2 mL of milliQ $H_2O$ via sonication. To prepare the Casp11 : KLA complexes, first, the desired amount of KLA was transferred to an Eppendorf tube and $H_2O$ was removed via SpeedVac. Subsequently, the KLA was resuspended in analysis buffer (see *SI Appendix,* Table S2), and sonicated. Notably, the analysis buffer pH was ≥ 7.4 to ensure that KLA exists in its physiological protonation state. The Casp11 protein of choice (in the same analysis buffer) was then mixed via pipetting and incubated at 25°C for 1 hour. At this point, the sample was ready for SEC-UV or CD experiments. For NMR, SEC-UV-RI-LS, and cryo-EM studies, the sample was additionally passed through a Superdex 200 Increase 10/300 GL column (Cytiva) equilibrated in SEC buffer (see *SI Appendix,* Table S2), to isolate the complexes from any unbound KLA micelles or other contaminants. The fractions containing the complexes were subsequently pooled and, if necessary, diluted to achieve the correct analysis buffer conditions. Finally, the sample was concentrated using a stirred cell concentrator with a 30-kDa MWCO, 25 mm diameter PES membrane.

***NMR measurements***

All NMR experiments were recorded at either 23.5 Tesla (1 GHz $^1$H frequency; Bruker Avance NEO), 18.8 Tesla (800 MHz; Bruker Avance III HD), 14.1 Tesla (600 MHz; Bruker Avance III HD), or 11.7 Tesla (500 MHz; Bruker Avance III HD) using spectrometers equipped with cryogenically cooled x, y, z axis pulsed-field gradient triple-resonance probes (23.5 T, 18.8 T, and 14.1 T) or a liquid nitrogen cooled z axis pulse-field gradient triple resonance probe (11.7 T). NMR data were processed using the NMRPipe suite of programs (8) and visualized using the Python3 nmrglue package (https://www.nmrglue.com/, (9)). Peak volumes were extracted using the Python3 peakipy package (https://j-brady.github.io/peakipy/).

*i) Assignment of crosspeaks* (*SI Appendix*, Fig. S3 and *SI Appendix*, Fig. S9)

The sequential backbone resonance assignment of the Casp11 CARD was achieved via a standard suite of triple-resonance experiments (HNCO, HN(CA)CO, HNCA, HN(CO)CA, HNCACB, CBCA(CO)NH, HNN, CCC-TOCSY (10, 11)) recorded at 14.1 T and -5°C, in either native or denaturing (6 M urea) conditions (See *SI Appendix, Table S2* for sample conditions). To obtain more complete assignments for the native condition, amide and carbonyl backbone resonances were mapped as a function of denaturant, using a series of HNCO spectra (12) recorded at the following urea concentrations: 0, 0.25, 0.5, 0.75, 1, 1.5, 2, 3, 4, 5, 6 M. The mapped chemical shifts of the amide and carbonyl resonances could then be used, in concert with HNCACB and CBCA(CO)NH experiments, to establish the Cα and Cβ chemical shifts for the corresponding residue. To minimize sample consumption, a sample-mixing approach was employed for the urea titration. In brief, two samples containing identical protein concentrations and either 0 M ("first" titration point) or 6 M ('last" titration point) urea were generated (See *SI Appendix, Table S2* for sample conditions), and HNCO spectra were obtained. These two samples were then mixed in a specific ratio so that two new samples containing the desired urea concentrations were obtained, as illustrated below. This was repeated until the mid-point of 3 M urea was reached:

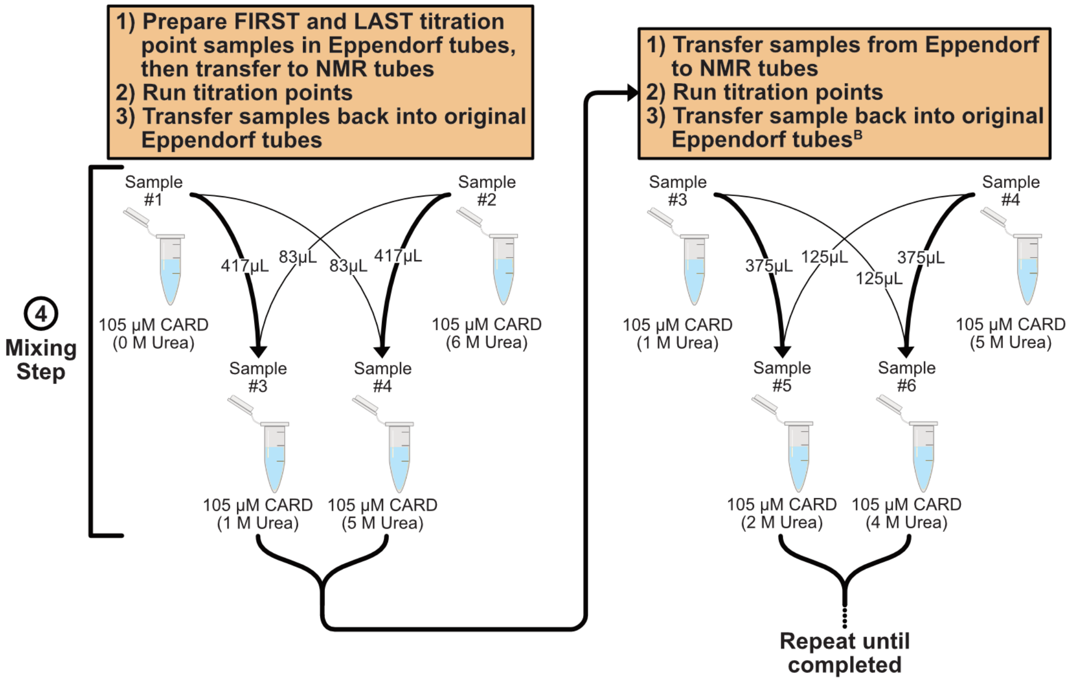


▲ *Example sample mixing scheme utilized for Casp11 CARD urea titration experiments. Note that, in practice, a small amount of sample is lost during every transfer step, so it is advisable to start with more than the required volume.*

All experiments utilized non-uniform sampling (NUS) with a Poisson-Gap sampling schedule, with sampling densities of between 20-35% (13). NUS-acquired datasets were reconstructed using the SMILE program in NMRPipe (14). Assignments were achieved using a combination of NMRFAM-SPARKY (15) and the I-PINE webserver (http://i-pine.nmrfam.wisc.edu/, (16)) software.

A significant number of the methyl sidechains of Met and Ile residues in the Casp11* PD variant were assigned via a mutagenesis strategy. In brief, 7/9 Met and 11/18 Ile residues were mutated (one at a time; typically Met → Leu and Ile → Met), and the resulting $^{1}H$-$^{13}C$ HSQC spectrum was compared to a "baseline" spectrum which contains no mutations. The disappearance of a resonance relative to the baseline can be attributed to the specific residue mutated. Additional assignments were obtained via 2D $^{13}C$-edited NOESY experiments recorded on a monomer mutant of the C254A Casp11 PD (R360E) that gave high quality spectra.

*ii) Extracting $k_{ex}$ and $p_B$ from $^{15}N$-CPMG measurements* (*SI Appendix*, Fig. S2B)

$^{15}N$-CPMG experiments were recorded on samples of Casp11 CARD with or without denaturant (*SI Appendix*, Table S2) at either 18.8 T (800 MHz, $^{1}H$ frequency) or 11.7 T (500 MHz, $^{1}H$ frequency) at -5°C using a constant-time CPMG pulse scheme in which $^{15}N$ magnetization is kept in-phase during the CPMG interval using a $^{1}H$ continuous-wave decoupling field (17). The constant-time relaxation interval, $T_{relax}$, was set to 30 ms with the frequency of chemical shift refocusing pulses, $\nu_{CPMG}$, ranging between 33 and 1000 Hz. A total of 17 values of $\nu_{CPMG}$ were acquired, along with two repeats for error estimation. Additionally, a reference plane without the constant-time relaxation interval was recorded, which provided reference intensity values ($I_0$)

allowing for calculation of $R_{2,eff} = -\frac{1}{T_{relax}} \cdot \ln\left(\frac{I}{I_0}\right)$, where $I$ is the intensity from the respective $\nu_{CPMG}$ plane.

Pseudo-3D CPMG experiments recorded on Casp11 CARD in buffer (native conditions) were analyzed using the ChemEx software package (https://github.com/gbouvignies/ChemEx). For all fits, the $R_1$ rates in both the ground and excited states were fixed to 2.6 $s^{-1}$ and 1.7 $s^{-1}$ for analysis of data recorded at 11.7 T and 18.8 T, respectively (based on calculated values according to the molecular weight of the CARD), while $^{15}N$ $R_2$ values for corresponding spins in ground and excited states were constrained to be the same. All residues were fit independently to a two-state model using data recorded at both 11.7 T and 18.8 T, so that per-residue values for exchange rates ($k_{ex}$) and minor state populations ($p_B$) were obtained. Experiments recorded on a sample in 6 M urea (denaturing conditions) were not analyzed as only flat dispersion profiles were present.

*iii) Diffusion measurements* (Fig. 3B)

A pulsed-field gradient diffusion scheme described previously (18), except with $^{15}N$ and $^{13}C$ pulses interchanged, was used to measure diffusion constants of Casp11 : KLA complexes from a series of 1D $^{13}C$-edited $^{1}H$ spectra (see *SI Appendix*, Table S2 for specific proteins and buffer compositions), 18.8 T and 25°C. A region of the $^{1}H$ signal extending between 2.1 to 1.9 ppm (Met residues) was integrated to obtain intensities as a function of gradient strength. A constant-time diffusion period of 200 ms was used for all measurements, along with dephasing and rephasing gradient lengths to ensure that approximately 40% of the initial signal remained in spectra recorded with the largest gradient strength. Translational diffusion constants could then be obtained using the relation:

$$I_{i,calc} = I_{0,calc} \times \exp(-A \cdot D \cdot G_i^2) \quad [S1]$$

where $I_{0,calc}$ and $I_{i,calc}$ are the calculated signal intensities in the absence and presence ($i = 1, \ldots, N$) of dephasing/rephasing gradients, D is the diffusion constant, $G_i$ is the gradient strength, and $A$ is a constant that depends on the experimental parameters. The difference between the calculated and experimental peak intensities ($I_{i,exp}$) could be fit via minimization of the residual sum-of-squared (RSS) values:

$$RSS = \sum_{i=1}^{N} \left[ I_{i,exp}\left(D, I_{0,exp}\right) - I_{i,calc}\left(D, I_{0,calc}\right) \right]^2 \qquad [S2]$$

using the Levenberg-Marquardt algorithm of the LMFIT (v1.3.4) Python software package (https://lmfit.github.io/lmfit-py/).

*iv) Correcting for differences in transverse relaxation rates of magnetization giving rise to monomer and dimer peaks* (Fig. 3C-D, and *SI Appendix*, Fig. S8A)

$^1$H transverse relaxation rates ($R_2$) for the monomer and dimer peaks were measured using a 2D $^1$H-$^{13}$C HSQC-based experiment that included a variable delay spin-echo element, with $^1$H relaxation rates extracted via fits of peak intensities to an exponential decay function. The extracted $R_2$ values for each monomer and dimer peak could then be used to correct peak volumes ($I_{exp}$) in spectra of Figures 3C,3D and *SI Appendix*, Figure S8A via the relation (19):

$$I_{corr} = I_{exp} \times \exp(R_2 \cdot \tau) \qquad [S3]$$

where $\tau = 0.0064$ s is the sum of the durations of the INEPT (20) and reverse INEPT transfers in the pulse scheme used. For the data presented in Figures 3C, 3D and S8A all titration points were corrected using $R_2$ values obtained at 260 µM (Casp11* PD), 60 µM (FL Casp11* + 3×KLA), and 830 µM (catalytically active D277A D285A Casp11 PD) protein concentrations, respectively.

*v) Determination of $K_D$ for Casp11* PD and catalytically active D277A D285A Casp11 PD* (Fig. 3C and *SI Appendix*, Fig. S8A)

Consider the dimerization reaction, $M + M \rightleftharpoons D$, where $M$ is the monomer and $D$ is the dimer, with the dissociation constant ($K_D$) given by:

$$K_D = \frac{[M]^2}{[D]} \qquad [S4]$$

Using the relation, $[T] = [M] + 2 \times [D]$, where $[T]$ is total protein concentration, the fraction dimer (fraction of monomeric units within a dimer), $Frac_D$, can be solved as:

$$Frac_{D_i}^{calc} = \frac{2 \times [D]}{[T]} \qquad [S5]$$

where

$$[D] = \frac{4 \times [T] + K_D - \sqrt{K_D^2 + 8 \times [T] \times K_D}}{8} \qquad [S6]$$

$Frac_{D_i}^{calc}$ values for each titration point can be calculated (using an initial guess of $K_D$) and compared to the experimental data $Frac_{D_i}^{expt}$,

$$Frac_{D_i}^{expt} = \frac{V_{D_i}}{V_{M_i} + V_{D_i}} \qquad [S7]$$

where $i = \text{M184, I250, M262}$, noting that the peak volumes for the monomer ($V_M$) and dimer ($V_D$) resonances are proportional to $[M]$, and $2 \times [D]$, respectively. Values of $K_D$ are extracted from minimization of the RSS differences between $Frac_{D_i}^{calc}$ and $Frac_{D_i}^{expt.}$

$$RSS = \sum_{j=1}^{N} \left[ Frac_{D_{i,j}}^{expt}([T]_j) - Frac_{D_{i,j}}^{calc}([T]_j, K_D) \right]^2 \qquad [S8]$$

using the Levenberg-Marquardt algorithm of the LMFIT (v1.3.4) Python software package.

*vi) Estimating effective concentration of Casp11* in the non-canonical inflammasome* (Fig. 3D)

The effective concentration of the Casp11* PDs is defined as the bulk concentration of Casp11* that would be required to achieve the same level of PD dimerization as for Casp11* in the non-canonical inflammasome. Starting from the relation:

$$K_D^{C11^*} = \frac{[M]^2}{[D]}, \qquad [S9]$$

it follows directly that

$$\frac{K_D^{C11^*}}{[M]_{eff}} = \frac{[M]_{eff}}{[D]_{eff}} = 2\frac{V_M}{V_D} \text{ or } [M]_{eff} = \frac{K_D^{C11^*} V_D}{2V_M} \qquad [S10]$$

where $\frac{V_M}{V_D}$ is the volume ratio of monomer – dimer pairs of peaks in spectra, corrected for differential relaxation during the fixed delays in the HSQC pulse sequence (see '*Correcting for differences in monomer-dimer peak transverse relaxation rates*'), and $[M]_{eff}$ and $[D]_{eff}$ are the effective concentrations of PD monomer and dimer that give rise to the fraction dimerization on the inflammasome. The effective concentration, $[T]_{eff}$, is then calculated as $[T]_{eff} = [M]_{eff} + \frac{2\times[M]_{eff}^2}{K_D^{C11^*}}$. Using the experimentally determined $K_D^{C11^*} (= 170 \pm 20$ µM) for the Casp11* PD (Fig. 3C) and the experimental *Frac*$_D$ (for example, at 10 $\mu$M bulk concentration of FL Casp11* *Frac*$_D$= $\frac{1}{1+\frac{V_M}{V_D}} = 0.75 \pm 0.02$) for the PD of FL Casp11* when bound to the non-canonical inflammasome complex (Fig. 3D), $[T]_{eff}^{10\mu M} = 1028 \pm 226$ µM. Note that *Frac*$_D$ is expected to be invariant to the bulk Casp11* concentration used, with values ranging from approximately 0.75 to 0.80 quantified in our experiments (Fig. 3D).

***Rationale for Casp11 mutations***

The mutations introduced in Casp11 can be grouped into three categories based on their function: (1) Resist protease degradation, (2) Eliminate spectral crowding around Met 262, which gives rise to intense monomer and dimer peaks whose intensities can be quantified accurately, and (3) increase PD dimerization affinity to improve quantification of the monomer–dimer equilibrium. In our hands, the C254A FL Casp11 was prone to proteolytic cleavage even after purification, likely due to trace *E. coli* protease contaminants. Unfortunately, commonly used protease inhibitors (PMSF, E-64, Pepstatin-A, EDTA, Bestatin, Leupeptin, Benzamidine) did not remedy this issue. Instead, we used an iterative process where sites of proteolysis were identified and then removed by mutagenesis. This was achieved by incubating 'purified' C254A FL Casp11 at 25°C for 48 hours (to simulate conditions during NMR experiments), followed by sample analysis using positive ion mode electrospray ionization mass spectrometry (ESI-MS). The accurate molecular weights of the products determined in this manner allowed identification of possible protease cleavage sites within the protein. Mutagenesis was performed to remove these tentative cut sites, and the process repeated until further cleavage products were not detected. The final set of mutations to alleviate protease cleavage were Q74N, T75N, F76L, F77L, L99A, T101S, L102A, L104S, C105A, S106P, P107H, L113A, C114S, L272A, C273S, producing the C254A FL Casp11$^{PR}$ variant (Fig 3B). Notably, while a handful of mutations were localized to the CARD, C254A FL Casp11$^{PR}$ retained its ability to interact with KLA and to form non-canonical inflammasome complexes, albeit with different populations of complexes I and III (*SI Appendix*, Fig. S7).

Due to the large size of the Casp11 : KLA complexes (ranging from approximately 200 - 450 kDa) and the low particle concentrations utilized, spectral sensitivity was critical to allow for reliable quantification of $Frac_D$. While both Ile 250 and Met 184 methyl probes gave rise to resolved peaks for monomer and dimer, their sensitivity was low and not amenable for studies of the complexes, especially since the FL protein could not be prepared with deuteration (poor expression yields). The intense (monomer and dimer) peaks from Met 262 overlapped with signals from other methionine crosspeaks. These Met residues were mutated to Gly (M1G), Ala (M92A), or Leu (M216L, M282L, M355L), alleviating the spectral crowding issue.

Finally, it was beneficial to introduce a mutation which increased the PD dimerization affinity, enlarging the dimer peak so that a more robust quantification of fraction dimer could be obtained. To achieve this a series of mutations at the dimer interface of Casp11 (using an X-ray structure of the dimer, PDBID: 6KMT (21)) were analyzed using the DDMut webserver (https://biosig.lab.uq.edu.au/ddmut/, (22)), and subsequent in-vitro analysis established that A361V shifted the equilibrium sufficiently (lowering $K_d$ by approximately 4-fold). The construct containing all three sets of mutations described above is referred to as FL Casp11*, or Casp11* PD with the exception that the PD version contains only mutations starting at residue 92 (Figure 3C-D).

***SEC-UV-RI-LS and SEC-UV measurements***

*i) SEC-UV-RI-LS* (Fig. 2A, and *SI Appendix*, Fig. S5)

All analytical SEC-UV-RI-LS measurements were performed using a Superdex 200 Increase 10/300 GL column connected to an Agilent 1260 Infinity II HPLC system with in-line UV-vis (set to 280 nm), Refractive Index (RI; Wyatt Optilab T-rEX) and Light Scattering (LS; Wyatt DAWN) detectors along with an autosampler. The system was equilibrated overnight at a flowrate of 0.1 mL/min with 20 mM NaPhos, 150 mM NaCl, 1 mM EDTA, pH 7.4 to ensure all detectors had reached a steady state. To normalize the LS detector, 60 μL of 2 mg/mL Bovine

Serum Albumin (BSA) was first injected and analyzed. For each run, 100 μL of sample containing either Casp11 CARD or CARD-fusion proteins prepared with a 3-fold molar equivalent of KLA (see '*Preparation of Casp11 : KLA complexes*') was injected and a flow rate of 0.30 or 0.35 mL/min was set so that each run lasted for a total of 88 or 72 min, respectively. Data was collected using the ASTRA software (v8.2), and was corrected for baseline, peak alignment, and band broadening prior to being exported for analysis using an in-house Python3 script.

Using the 'three-detector' method (23), the total molecular weight of the complex, along with the molecular weights of the proteinaceous and KLA components were elucidated. This method relies on knowing the extinction coefficient ($\varepsilon_{280}$, in this case at 280 nm) and the refractive index increment ($dn/dc$) for each component. For the Casp11 constructs used, the $\varepsilon_{280}$ values (in units of L $g^{-1}$ $cm^{-1}$; estimated using ExPASY ProtParam, https://web.expasy.org/protparam/) and $dn/dc$ values (in units of mL/g; estimated using SEDFIT v18.1, https://sedfitsedphat.nibib.nih.gov/software, (24)) are given in *SI Appendix*, Table S1. KLA has no absorbance at 280 nm (thus $\varepsilon_{280} = 0$) and as an exact $dn/dc$ value is not known, we used a range between 0.140 to 0.160 (based on $dn/dc$ values for other lipids) in our analysis (25, 26). The approach followed is based on a description by Slotboom and coworkers (23) and summarized here.

For each of complexes I, II, and III (Fig. 3A) we first calculated *δ*, the amount of KLA associated with protein (g/g), according to:

$$\frac{1}{1+\delta}(dn/dc)_{Casp11} + \frac{\delta}{1+\delta}(dn/dc)_{KLA} = \frac{\Delta RI}{\Delta UV_{280}}\left(\left(\frac{1}{1+\delta}\right)\varepsilon_{280}\right) \qquad [S11]$$

where $\Delta RI$ and $\Delta UV_{280}$ are excess signal (relative to buffer) from the RI and UV-vis detectors at a given timepoint in the SEC trace, respectively. Using the $\delta$ values for each complex, the apparent $dn/dc$ value of the complex can be calculated according to:

$$dn/dc_{complex} = K_1 \cdot \frac{\Delta RI}{\Delta UV_{280}} \left( \left( \frac{1}{1+\delta} \right) \varepsilon_{280} \right) \qquad [S12]$$

where $K_1$ is a calibration factor that is $\approx 1$ (0.97-1.01 in our case). The mass concentration of the complex ($c_{complex}$) at each timepoint in the SEC trace can be calculated using:

$$c_{complex} = \frac{\Delta UV_{280} \times (1+\delta)}{\varepsilon_{280}} \quad . \qquad [S13]$$

From Eqs. [*S12*] and [*S13*] the total molecular weight of the complex ($MW_{complex}$) is obtained from:

$$MW_{complex} = K_2 \cdot \frac{\Delta LS}{\left(dn/dc_{complex}\right)^2 \times c_{complex}} \qquad [S14]$$

where $\Delta LS$ is the excess signal (over buffer) from the light-scattering detector and $K_2$ is a constant that depends, among other things, on the wavelength of light used and the angle between the incident and scattered light (23). Here we have used 15 different scattering angles so that the average MW of the complex across all angles was obtained. Finally, the molecular weights of the Casp11 ($MW_{Casp11}$) and KLA ($MW_{KLA}$) components were obtained from:

$$MW_{Casp11} = \frac{MW_{complex}}{1+\delta} \qquad [S15]$$

and

$$MW_{KLA} = MW_{complex} - MW_{Casp11} \,. \qquad [S16]$$

The stoichiometric ratios could then be easily determined, using the theoretical molecular weights of the Casp11 (see *SI Appendix,* Table S1; determined via ExPASY ProtParam) and KLA (2306.84 g/mol) components.

*ii) SEC* (Fig. 3A)

All analytical SEC measurements were performed using a Superdex 200 Increase 10/300 GL column connected to an Agilent 1260 Infinity II HPLC system with a UV-vis detector (set to 280 nm) and an autosampler. The system was equilibrated with 20 mM NaPhos, 150 mM NaCl, 1 mM EDTA, pH 7.4. For each run, 100 μL of sample containing either 50 μM (replicate 1) or 35 μM (replicate 2) C254A FL Casp11 with between 0 to 5-fold molar equivalents of KLA (see '*Preparation of Casp11 : KLA complexes*') was injected, with the flowrate set to 0.4 mL/min and each run lasting 75 min.

Due to overlap of complexes I-III in the SEC trace, a deconvolution procedure was required to obtain the fractional populations of each (Fig. 3A, top). This was achieved via an in-house Python3 script which modelled each complex peak as a Gaussian function:

$$f^{g}(x) = A \times \exp\left(\frac{-(x-\mu)^2}{2\sigma^2}\right), \qquad [S17]$$

where $A$ is the amplitude, $\mu$ is the center, and $\sigma$ is width of the peak. In addition, the unbound Casp11 peak was modelled as an exponential Gaussian:

$$f^{eg}(x) = \frac{A\gamma}{2} \times \exp\left(\gamma\left(\mu - x + \frac{\gamma\sigma^2}{2}\right)\right)\text{erfc}\left(\frac{\mu + \gamma\sigma^2 - x}{\sqrt{2}\sigma}\right), \qquad [S18]$$

where $\gamma$ is the rate of the exponential component, or skew (27, 28). Two additional Gaussian functions along with one additional exponential Gaussian function were included to account for contaminants and the void, respectively. The sum of these Gaussian and exponential Gaussian curves could be fit to the experimental SEC traces using a least-squares fitting procedure, minimizing RSS (using the Levenberg-Marquardt algorithm in LMFIT 1.3.4), the difference between the experimental ($f^{expt}(x)$) and calculated ($f_{SUM}^{calc}(x)$) SEC traces at each KLA/Casp11 value (referred to as KLA ratio; $j = 0, 0.25, 0.5, 1, 2, 3, 5$):

$$f_{SUM}^{calc}(x) = f_{I}^{g}(x) + f_{II}^{g}(x) + f_{III}^{g}(x) + f_{Free}^{eg}(x) + C(x) \qquad [S19]$$

$$RSS = \sum_{j} \left[ f_{j}^{expt}(x) - f_{SUM,j}^{calc}(x) \right]^2 \qquad [S20]$$

In Eq. $[S19]$ $f_{I-III}^{g}(x)$ are the Gaussian peaks for each complex I-III, $f_{Free}^{eg}(x)$ is the exponential Gaussian peak for the unbound Casp11, and $C(x)$ is the sum of the additional Gaussian and exponential Gaussians associated with contaminants and void. The center ($\mu$), width ($\sigma$), and skew ($\gamma$, for exponential Gaussians only) were fit as single values on a per-peak basis across all KLA ratios. In contrast, the amplitude ($A$) of each peak was fit to each unique KLA ratio.

The data present at the bottom of Figure 3A were calculated using the amplitude values for each peak to determine the fraction of each species ($i = I, II, III, Free$) as a function of the KLA ratio. Here, the contributions from the contaminants and void were omitted:

$$Frac_{i,j} = \frac{A_{i,j}}{A_{I,j} + A_{II,j} + A_{III,j} + A_{Free,j}} \qquad [S21]$$

***Circular Dichroism (CD) measurements*** (*SI Appendix*, Fig. S4)

All CD measurements were performed on a Jasco J-1500 CD spectrophotometer at 25ºC using 1 mm quartz cuvettes. Spectra were acquired between 260-185 nm using 0.2 nm steps and were baseline corrected against a buffer-only sample. Exact sample composition is described in *SI Appendix*, Table S2, but importantly contained NaF in place of NaCl to allow for reliable data acquisition below 200 nm. Secondary structure components were deconvoluted using the BeStSel webserver Single Spectrum Analysis function (https://bestsel.elte.hu/, (29, 30)) with similar results being obtained using the SELCON2/3 (31) and SESCA (32) algorithms in the ChiraKit webserver (https://spc.embl-hamburg.de/app/chirakit, (33)).

***Cryo-EM and image analysis*** (*SI Appendix*, Fig. S6)

A 3.6 mg/mL (85 µM) sample of C254A FL Casp11 + 3×KLA (see *SI Appendix*, Table S2 for sample conditions) was applied to homemade nanofabricated holey gold grids with regular arrays of ~2 µm holes (34). Grids were glow-discharged in air for 2 min, and 1.5 µL of sample was applied to each grid, blotted for 2 s at 4 °C and 80% relative humidity, and frozen in liquid ethane with a Leica EM GP2 freezing device. Samples were imaged with a Thermo Fisher Scientific Glacios electron microscope operating at 200 kV and equipped with a Falcon 4i camera. Data collection was automated with the *EPU* software package (Thermo Fisher Scientific). Electron event representation movies were collected (35) with a calibrated pixel size of 1.5 Å/pixel and a total exposure of ~40 $e^-/Å^2$, and fractionated into 30 exposure fractions.

Movie alignment with patch-based motion correction and estimation of contrast transfer function (CTF) parameters were performed with *cryoSPARC Live* (36). All other image analysis steps were performed with *cryoSPARC*. After removing movies with undesirable CTF fit, ice thickness, or motion, 260 movies were selected for further processing. Templates for particle selection were generated from 2D classification of manually selected particle images. The dataset was cleaned with multiple rounds of 2D classification, providing 215,396 particle images. While micrographs showed well-dispersed particles and 2D classification led to consistent class averages that were ~100 Å in diameter, 3D reconstruction failed to produce a coherent 3D structure. The inability to calculate a 3D map from the particle images is likely the result of conformational and compositional heterogeneity within the particle population, arising from the flexible linker between domains and variability in oligomeric assembly. These heterogeneous particle images result in over-refinement of heterogeneous features in 2D class average images and prevent identification of a homogeneous subset of images for high-resolution reconstruction.

***AUC measurements*** (*SI Appendix*, Fig. S8B)

*SV-AUC Experiments*

Sedimentation velocity (SV) analytical ultracentrifugation (AUC) experiments were carried out using a Beckman Coulter ProteomeLab XL-I ultracentrifuge following a previously established protocol (37). In brief, samples of wildtype (WT) Casp11 PD (see *SI Appendix*, Table S2 for sample conditions) were loaded into AUC cell assemblies equipped with either 12 mm or 3 mm charcoal-filled Epon double-sector centerpieces. These were then inserted into an An-50 Ti analytical rotor, followed by temperature equilibration in the centrifuge at 25°C for 2-3 hrs. After accelerating the rotor to 50 krpm, radial scans of each sample were collected using absorbance (230 nm) optical detection systems.

*Sedimentation Velocity (SV) Data Analysis*

Analysis of SV data recorded on WT Casp11 PD was performed in SEDFIT (V18.1) using the standard c(s) model (38). The c(s) distributions were normalized to the standard condition (water at 20°C) in GUSSI (39) using the calculated buffer density and viscosity values obtained from SEDNTERP (40). To generate an isotherm of the protein concentration series, weighted sedimentation values, $s_{20,\mathrm{w}}$, were obtained from integration of c(s) distributions in GUSSI. The resulting $s_{20,\mathrm{w}}$ isotherm was analyzed in SEDPHAT, and fit to a monomer-dimer self-association model (37), allowing for the dissociation constant ($K_D^{mature}$) to be extracted. While the s-values for the monomer (2.74 S) and dimer (4.11 S) are well defined based on their peak positions in the c(s) distributions, due to the presence of a contaminant sedimenting around 2 S (~ 10% of total signal), the $s_{20,\mathrm{w}}$ values obtained via integration of the c(s) distributions are systematically shifted. To account for this, 'effective' monomer ($s_{\mathrm{M}}$) and dimer ($s_{\mathrm{D}}$) sedimentation coefficients were estimated and constrained to [2.56, 2.83] S and [3.79, 3.98] S, respectively in the analysis of the $s_{20,\mathrm{w}}$ isotherm.

Supporting Information Figures

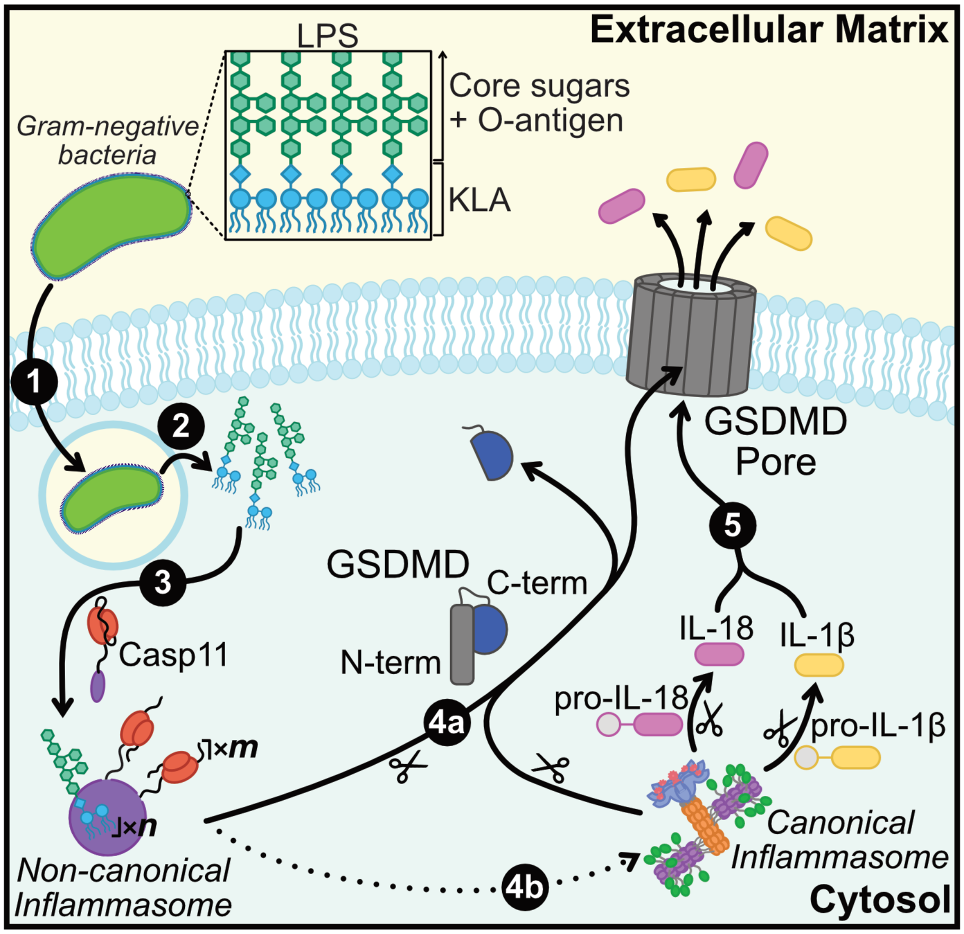


**Figure S1.** Schematic of the inflammatory cell death pathway highlighting the roles of both the non-canonical and canonical inflammasomes (41). In brief, gram-negative bacteria, which contain an outer membrane composed of lipopolysaccharide (LPS), are phagocytosed by immune cells **(1)** which eventually leads to release of their LPS into the cytosol **(2)**. The cytosolic LPS binds to the Casp11 CARD, triggering the formation of the non-canonical inflammasome complex composed of $m$ copies of Casp11 and $n$ copies of LPS **(3)** and upregulating Casp11 proteolytic activity. Activated Casp11 cleaves Gasdermin-D (GSDMD), releasing the pore-forming N-terminal domain from the inhibitory C-terminal domain **(4a)**. Further downstream processes can also lead to the formation of the canonical inflammasome (42–44), which also cleaves GSDMD along with prointerleukin1β (pro-IL-1β) and pro-IL-18, forming mature IL-1β and IL-18, respectively **(4b)**. IL-18 and IL-1β can then pass-through the plasma-membrane embedded GSDMD pore **(5)**, entering the extracellular matrix where they can bind to surface receptors on neighbouring cells, propagating the inflammatory response.

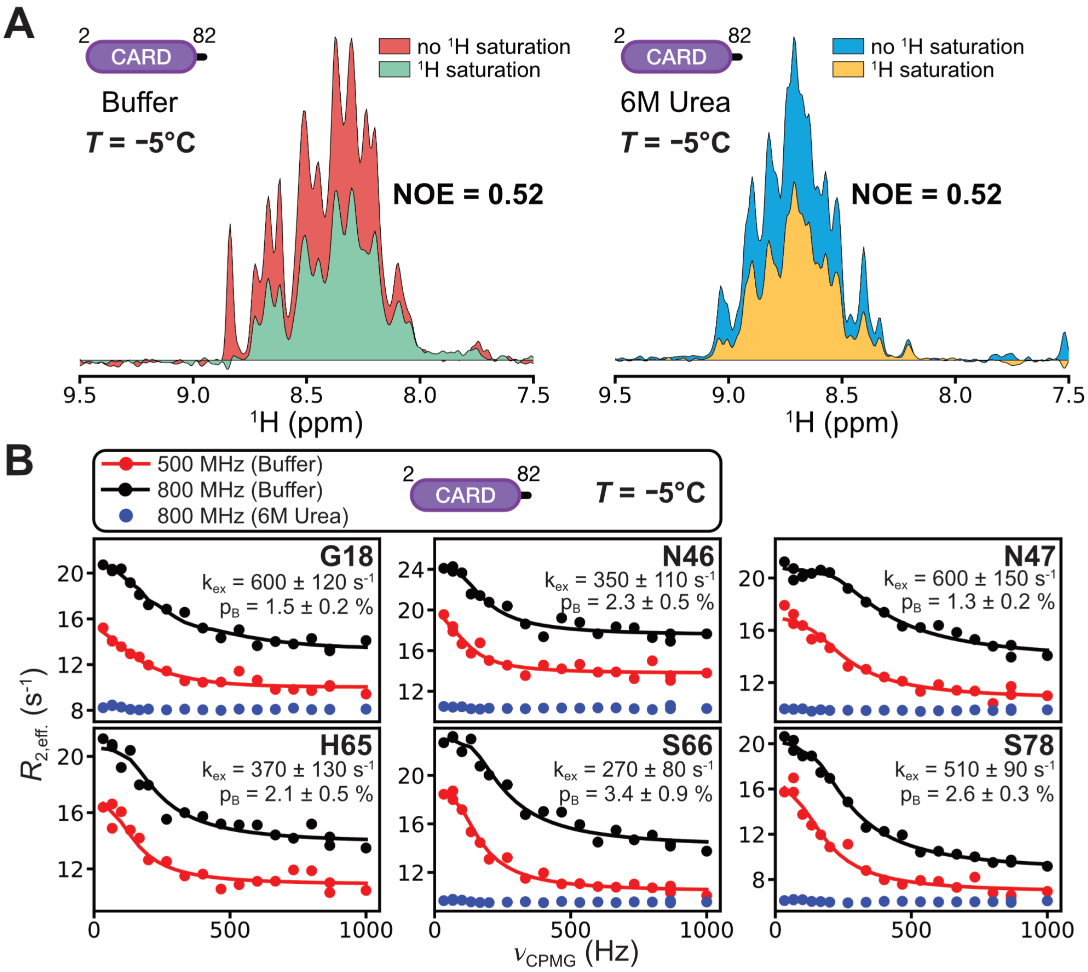


**Figure S2. (A)** 1D $^{15}$N{$^{1}$H} hetNOE spectra of the Casp11 CARD in buffer (left) and in 6 M urea (right) recorded at 18.1 T (800 MHz, $^{1}$H frequency) and -5°C. The average NOE value, calculated as the area under the '$^{1}$H saturation' spectrum divided by the area under the 'no $^{1}$H saturation' spectrum. **(B)** Experimental $^{15}$N-CPMG dispersion profiles for select Casp11 CARD residues recorded at either 500 MHz ($^{1}$H frequency, red circles) or 800 MHz ($^{1}$H frequency, black circles) in buffer at -5°C were each independently fit to a two-state model (solid lines) using the ChemEx program (https://github.com/gbouvignies/chemex). Extracted per-residue exchange rates ($k_{ex}$) and excited state populations ($p_B$) are shown. Addition of 6 M Urea (blue; 800 MHz, $^{1}$H frequency) results in flat dispersion profiles due to quenching of dynamics on the μs-ms timescale. Residue H65 was overlapped in 6 M urea preventing analysis.

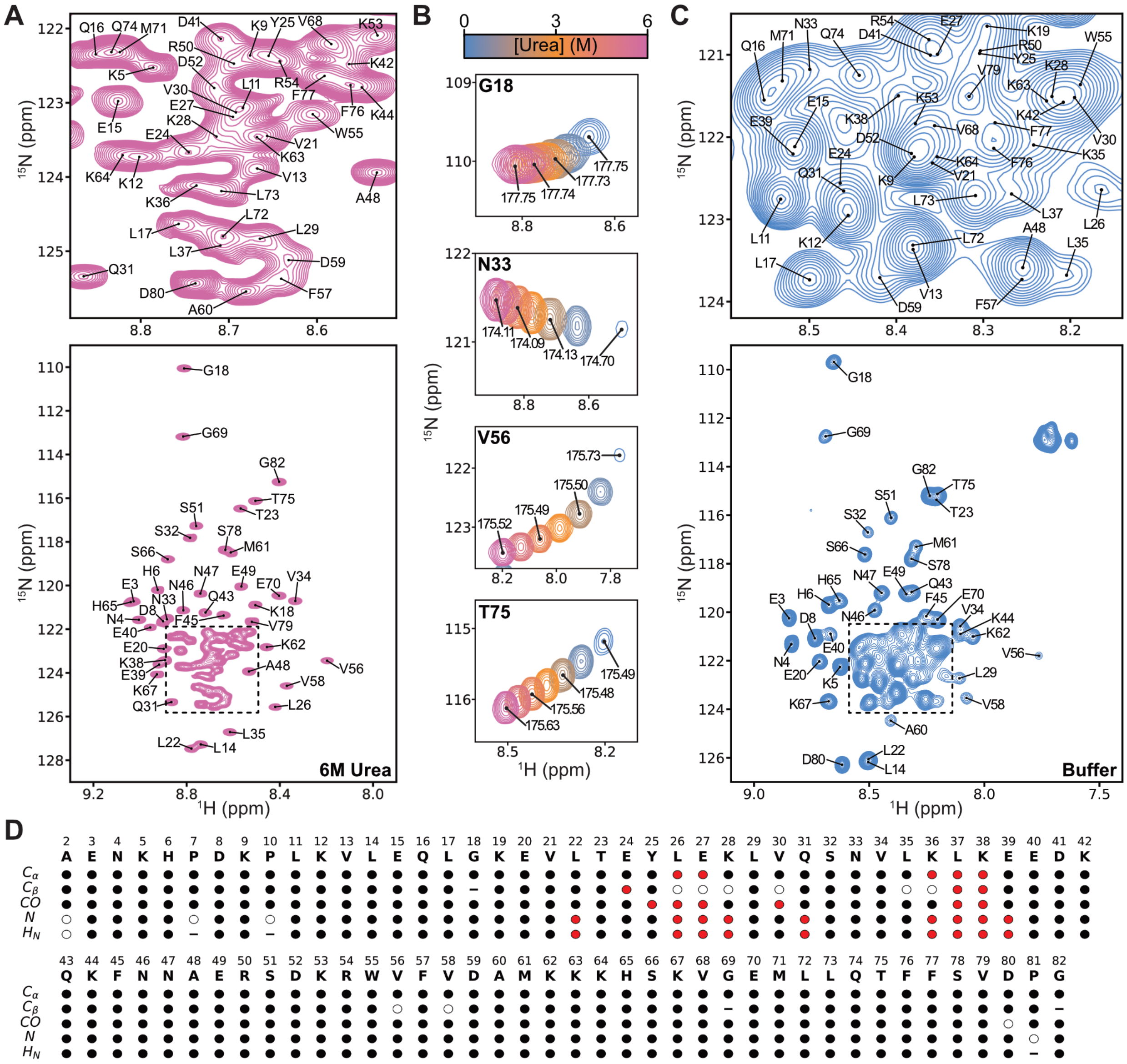

**Figure S3.** Assignment of the Casp11 CARD. **(A)** 2D $^{1}H$-$^{15}N$ HSQC spectrum of U-$^{13}C$,$^{15}N$ Casp11 CARD in 6 M urea recorded at 14.1 T (600 MHz, $^{1}H$ frequency) and -5°C. Assignments for all amide resonances are indicated, with the top panel an enlarged view of the region in the dashed-box (bottom). **(B)** Urea titration profiles for select residues, highlighting the large change in amide peak positions as the concentration of urea varies from 0 M to 6 M. Carbonyl chemical shifts, which are much less sensitive to changes induced by urea, are annotated for each residue. Using this approach, the chemical shift assignments in 6 M urea could be transferred to the 0 M urea condition. **(C)** 2D $^1$H-$^{15}$N HSQC spectrum of U-$^{13}$C,$^{15}$N Casp11 CARD in buffer recorded at 14.1 T (600 MHz, $^1$H frequency) and -5°C. Assignments for all amide resonances are indicated, with the top panel showing an enlarged view of the region in the dashed-box (bottom). **(D)** Sequence coverage of Casp11 CARD assignments. Filled black circles indicate that the resonance could be assigned in buffer, while filled red circles indicate that the assignment was only achieved via the urea titration. Unfilled circles and '–' indicate no assignment or not applicable ($C_\beta$ for Gly, $H_N$ for Pro), respectively.

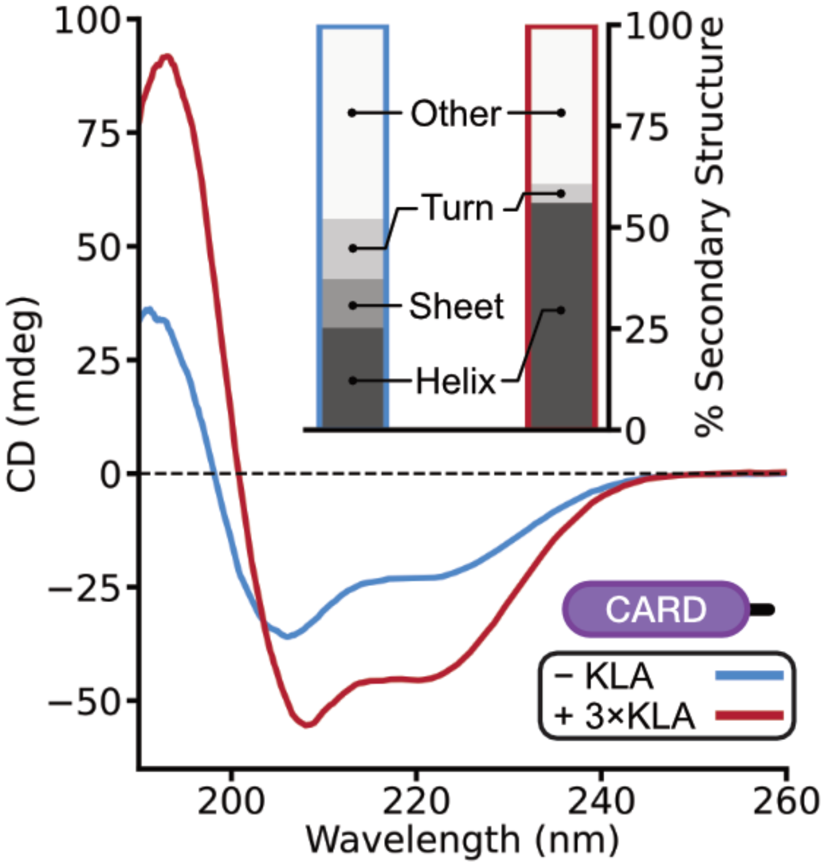


**Figure S4.** Circular dichroism measurements of Casp11 CARD in the absence (blue trace) and presence of 3-fold molar equivalent KLA (red trace), 25°C. The percent fraction of the indicated secondary structure element was determined (inset) using the BeStSel webserver (https://bestsel.elte.hu/, (29, 30)). In the absence of KLA, we obtain 25% α-helix, 12% β-sheet, 15% turn, and 48% 'other'. In the presence of 3-fold molar equivalent KLA, we obtain 56% α-helix, 0% β-sheet, 5% turn, and 39% 'other'.

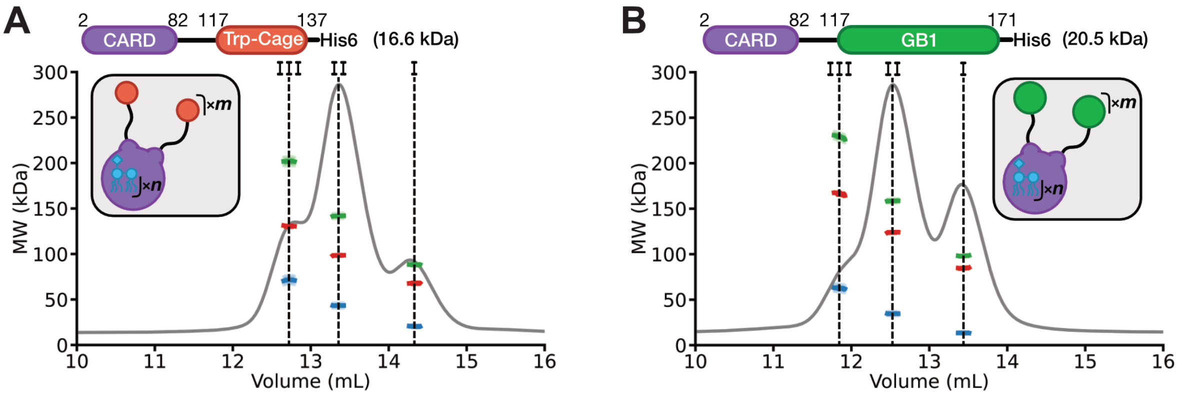


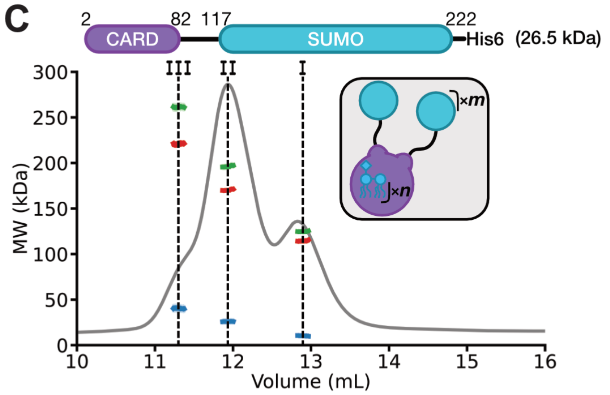


**D**

| | I | | II | | III | |
|---|---|---|---|---|---|---|
| | Protein | KLA | Protein | KLA | Protein | KLA |
| CARD-TrpCage | 4.2 ± 0.2 | 9.6 ± 0.9 | 5.9 ± 0.1 | 19.3 ± 1.9 | 7.8 ± 0.2 | 31.0 ± 3.0 |
| CARD-GB1 | 4.0 ± 0.2 | 5.6 ± 0.6 | 5.9 ± 0.2 | 14.5 ± 1.4 | 7.9 ± 0.3 | 27.4 ± 2.6 |
| CARD-SUMO | 4.5 ± 0.3 | 4.2 ± 0.4 | 6.6 ± 0.3 | 11.7 ± 1.1 | 8.6 ± 0.5 | 18.1 ± 1.7 |

**Figure S5.** SEC-MALS data recorded on mimics of the non-canonical inflammasome where the PD of Casp11 is replaced by **(A)** Trp-Cage (30 amino acids), **(B)** GB1 (64 amino acids), and **(C)** SUMO (115 amino acids) in the presence of 3-fold molar equivalent KLA. Domain organization of each construct and their corresponding monomeric molecular weight are shown at the top of each panel, along with a cartoon of each complex, consisting of *m* protein protomers and *n* KLA molecules in the grey boxes. For each construct, a UV-trace (grey; y-axis not shown) indicates that three unique complexes can be separated (annotated as I, II, and III; dotted black lines). Using the 'three-detector' method (See *SI Appendix Text*, (23)), the molecular weight of the complex (green), as well as the protein (red) and KLA (blue) components can be elucidated. Errors from the uncertainty in the change in refractive index with KLA concentration are illustrated with lighter-shade bars for the complex and KLA. **(D)** The protein and KLA stoichiometries for each construct are tabulated, with errors calculated using n=2 experimental replicates.

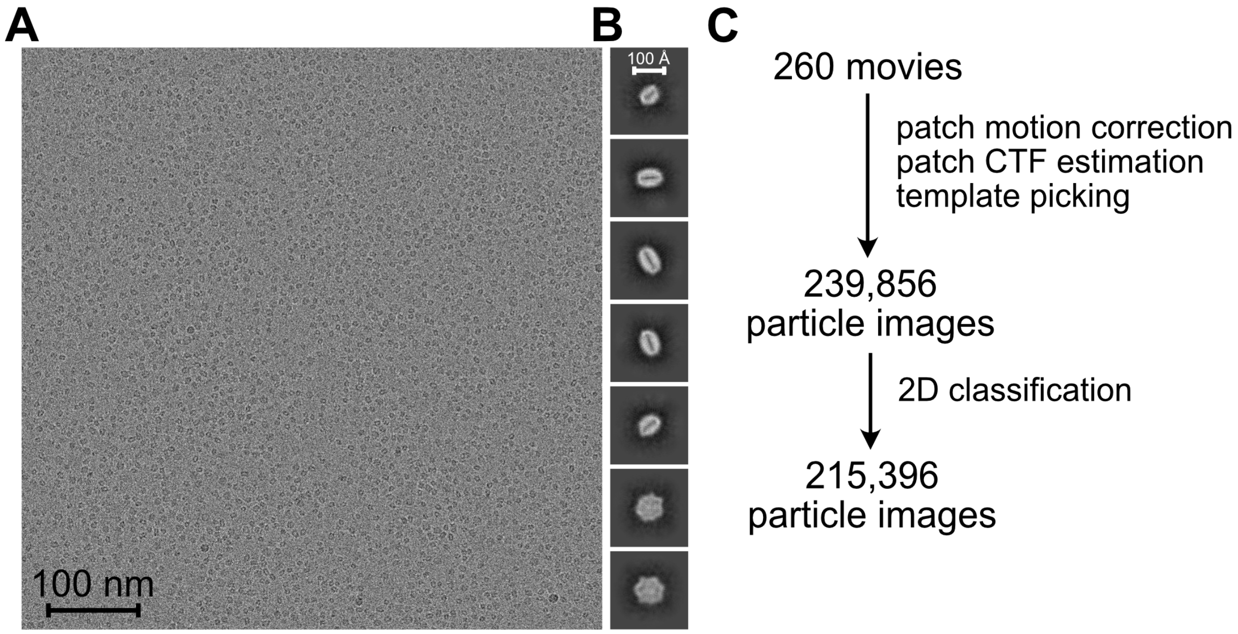


**Figure S6. (A)** Representative micrograph from cryo-EM data collection of a 3.6 mg/mL sample of SEC-filtered C254A FL Casp11 with 3-fold molar excess KLA and **(B)** 2D class average images. **(C)** Workflow employed to obtain 2D class averages. As discussed above, it was not possible to identify a homogeneous subset of images for high-resolution reconstruction.

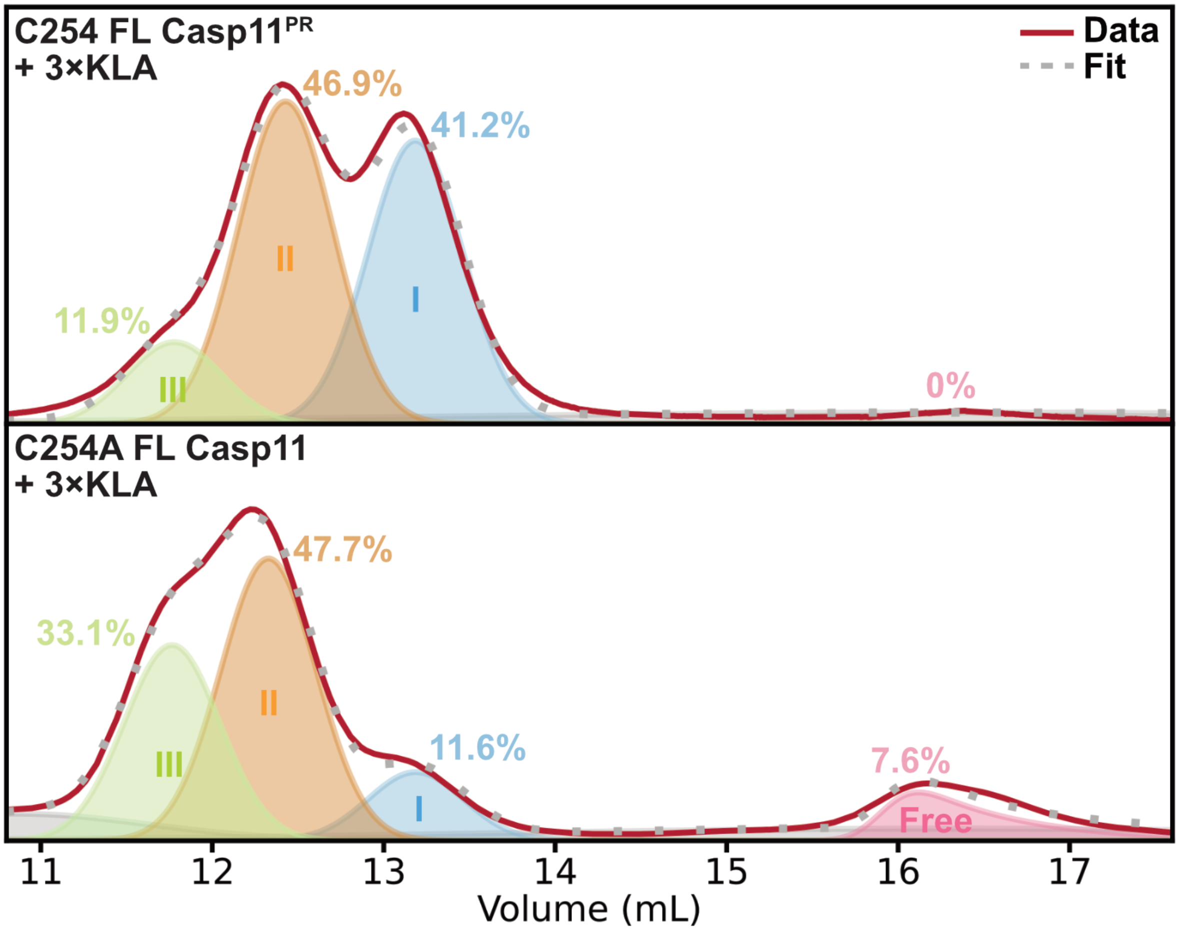


**Figure S7.** Comparison of size-exclusion profiles for C254A FL Casp11$^{PR}$ (top) and a representative replicate of C254A FL Casp11 (bottom) in the presence of 3-fold molar excess KLA (red line). Each peak is fit to either a Gaussian (I: blue, II: orange, III: green) or a skewed-Gaussian (Free: pink) lineshape. Two additional Gaussian and one additional skewed-Gaussian functions are included in the fit to account for contaminants/aggregates (grey), with the summation of all fitted components represented by a dotted grey line in each panel. The fraction of each fitted component is annotated next to the respective peak.

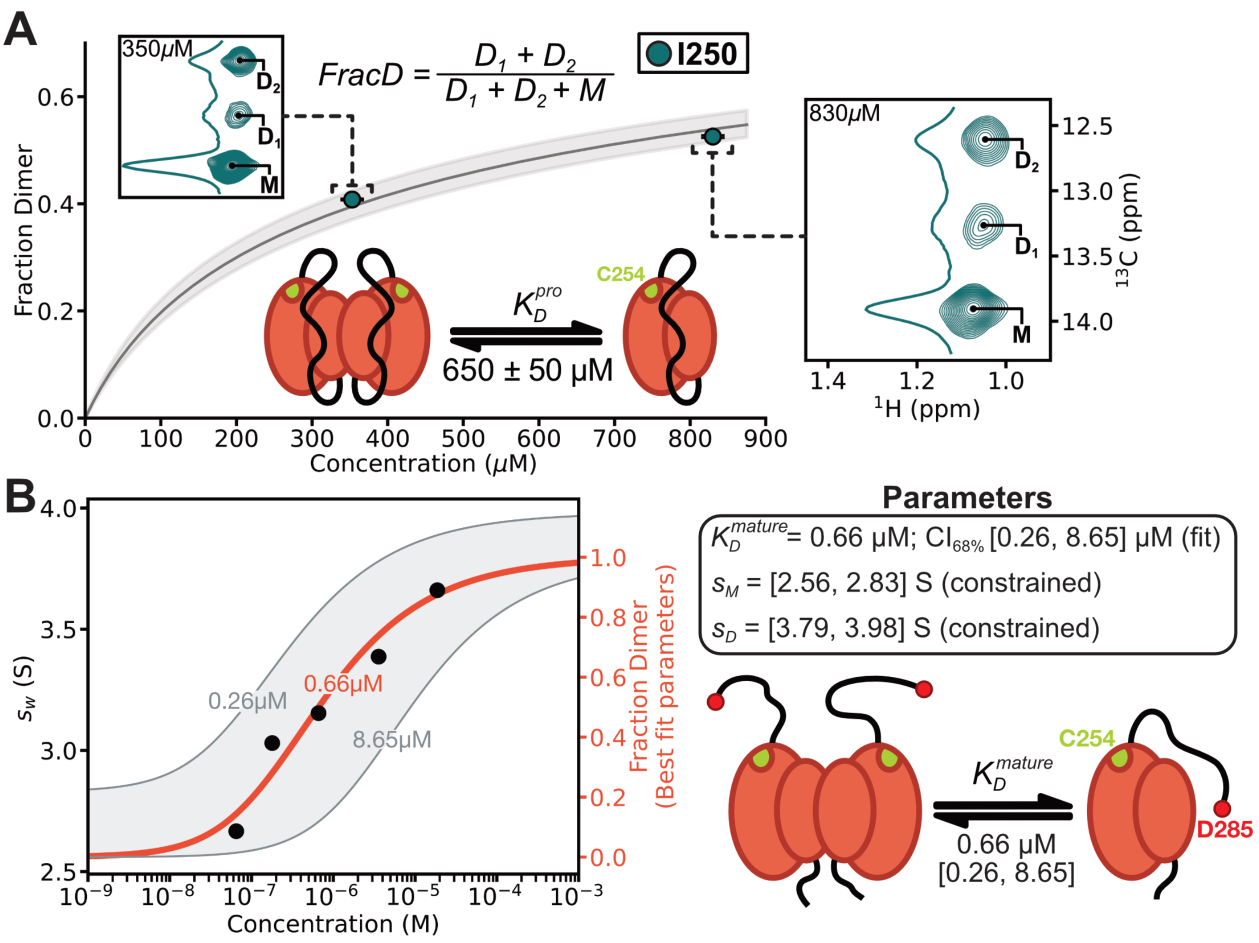


**Figure S8. (A)** Fraction dimer quantified from 2D $^{1}H$-$^{13}C$ HSQC spectra using a $^{13}C$-Ileδ1 labelled catalytically active D277A D285A Casp11 PD (protonated) sample, quantified from the I250 peak volumes $\left(Frac_D = \frac{D_1+D_2}{M+D_1+D_2}\right)$, and corrected for differences in relaxation rates between $^{1}H$ magnetization derived from monomers and dimers (filled circles). Notably, the spectrum of the catalytically active protein shows three peaks for the I250 δ1 methyl group, with two from the dimer state and a single peak derived from the monomer. The identities of these peaks were confirmed via a "monomer mutant" (R360E) for which only the monomer peak was observed, a I250L mutant which resulted in all three peaks disappearing, and by recording spectra at two different concentrations (350 µM, left inset and 830 µM, right inset) and quantifying normalized peak intensities. Notably, at higher concentration the normalized monomer peak volume decreased

($M^{830}/M^{350}$ = 0.77) and the two normalized dimer peak volumes increased ($D_1^{830}/D_1^{350} = 1.33$, $D_2^{830}/D_2^{350} = 1.17$), as expected. The experimental fraction dimer was fit to a monomer-dimer dissociation model ($M + M \rightleftharpoons D$), from which a dimer dissociation constant ($K_D^{pro}$) was determined (650 ± 50 μM). The data were recorded at 14.1 T (600 MHz, $^1$H frequency) and 25°C. A cartoon illustrating the monomer-dimer equilibrium of catalytically active D277A D285A Casp11 PD is displayed. The Asp to Ala mutations preclude self-cleavage of the p10-p20 linker which would normally accompany dimer formation. **(B)** Weighted sedimentation values, $s_w$, as a function of the concentration of WT Casp11 PD, measured by sedimentation velocity AUC at 25°C (black circles). $s_w$ values were fit to a monomer-dimer dissociation model ($M + M \rightleftharpoons D$) with monomer ($s_M$) and dimer ($s_D$) sedimentation coefficients constrained to [2.56, 2.83] S and [3.79, 3.98] S, respectively. The best fit $K_D^{mature}$ value was determined to be 0.66 μM (thick orange line), with a 68% confidence interval ($CI_{68\%}$) of $0.26\ \mu M < K_D^{mature} \leq 8.65\ \mu M$ (thin grey lines represent limits with shading showing the $CI_{68\%}$). The best fit $K_D^{mature}$, $s_M$ and $s_D$ values were used to calculate the fraction dimer profile (right y-axis, orange) as a function of concentration. A cartoon illustrating the monomer-dimer equilibrium of WT Casp11 PD is displayed in the bottom left, showing the cleavage at D285 that occurs in the WT protein.

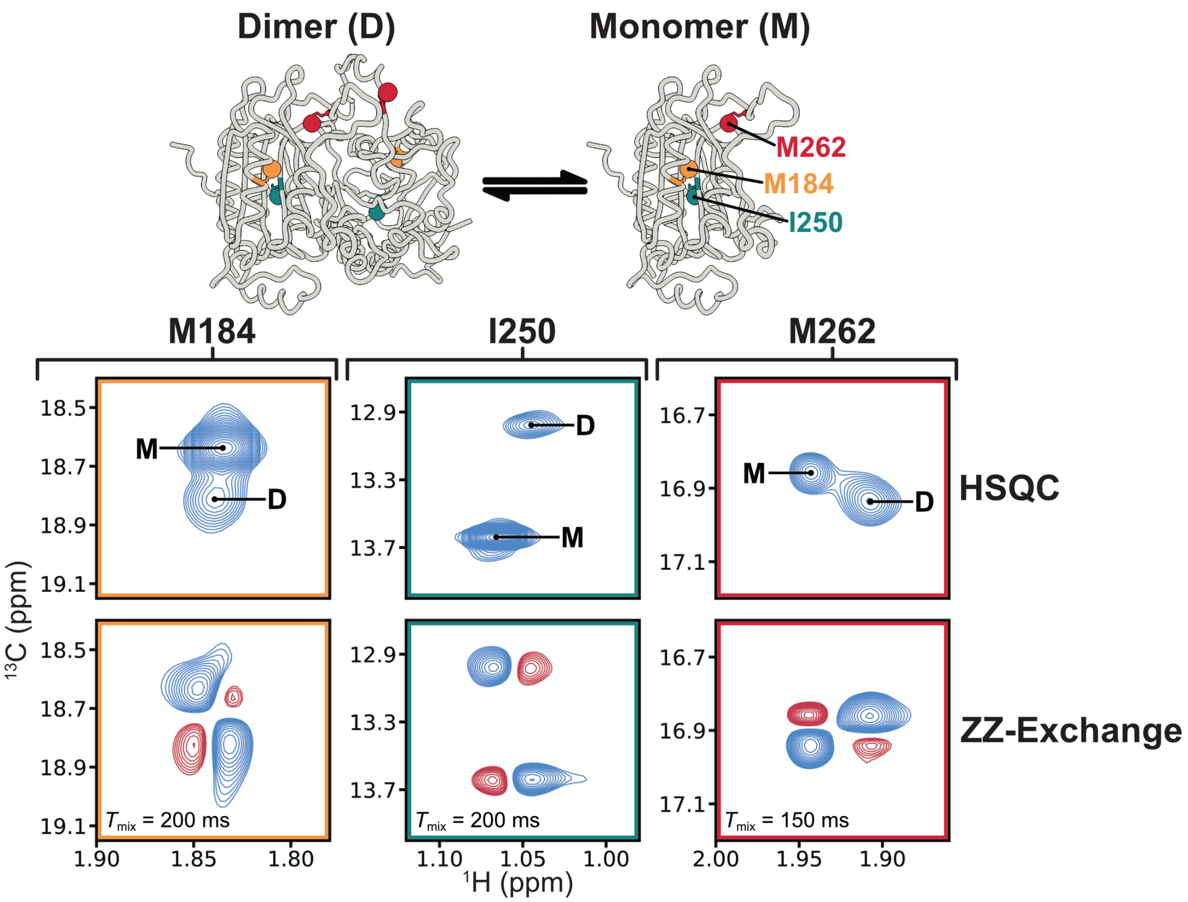


**Figure S9.** Methyl group probes in Casp11* PD with unique monomer-dimer chemical shifts in 2D $^{1}$H-$^{13}$C HSQC spectra were used to measure $K_D^{C11^*}$. The corresponding peaks were assigned via mutagenesis (see *SI Appendix Text*, '*Assignment of crosspeaks*') and by 2D $^{1}$H-$^{13}$C ZZ-exchange experiments recorded using either U-$^{2}$H, ILVM-labelled C254A Casp11 PD (M184 and I250; $T_{mix}$ = 200 ms) or U-$^{1}$H, ILVM-labelled Casp11* PD (M262; $T_{mix}$ = 150 ms). The difference between a pair of spectra where the $^{13}$C chemical shift was acquired either before or after the $T_{mix}$ period (during which exchange occurs) highlights the presence of "COSY-type" cross-peaks that are indicative of monomer-dimer interconversion on a timescale of many tens to hundreds of milliseconds. Experiments were recorded at 23.5 T (1 GHz, $^{1}$H frequency) and 25°C. A cartoon representation (45) of the Casp11 PD AlphaFold structure (AF-P70343-F1-v6, (46, 47)) is shown above, highlighting the location of the methyl group probes.

**Table S1.** DNA and amino acid sequences for select constructs.

| Construct | Sequence | Notes |
|---|---|---|
| **Casp11 CARD** | **DNA:**<br>ATGGCTGAAAATAAACATCCGGATAAGCCTCTCAAAGTTCTGGAACAATTAGGGAAG<br>GAAGTCCTGACCGAATACCTGGAAAAACTTGTACAAAGTAATGTACTCAAATTGAAG<br>GAAGAAGACAAACAAAAGTTTAATAATGCAGAGCGTTCTGACAAGCGTTGGGTCTTC<br>GTGGACGCGATGAAGAAGAAGCATAGTAAGGTTGGTGAAATGCTCCTTCAGACATT<br>CTTTAGTGTGGATCCGGGTTCGCATCATTGGTCTATTAATCACCATCACCACCATCAT<br>TCTGCATGGAGCCATCCTCAGTTTGAAAAGGGTGGCGGGAGCGGTGGTGGTTCCG<br>GCGGGTCTGCCTGGTCCCATCCTCAATTCGAGAAGTAA<br>**Amino Acids:**<br>MAENKHPDKPLKVLEQLGKEVLTEYLEKLVQSNVLKLKEEDKQKFNNAERSDKRWVF<br>VDAMKKKHSKVGEMLLQTFFSVDPG↓SHHWSINHHHHHHSAWSHPQFEKGGGS<br>GGGSGGSAWSHPQFEK* | ↓ = SNAC cut-site<br>MW = 9,457 Da<br>$\varepsilon_{280}$ = 6,990 $M^{-1}$ $cm^{-1}$<br>= 0.739 L $g^{-1}$ $cm^{-1}$<br>dn/dc = 0.187 mL/g |
| **C254A FL Casp11** | **DNA:**<br>ATGAAACACCACCACCACCACCACCCGATGAGTACCGAAAATCTGTACTTCCAGG<br>GCATGGCTGAAAATAAACATCCGGATAAGCCTCTCAAAGTTCTGGAACAATTAGGGA<br>AGGAAGTCCTGACCGAATACCTGGAAAAACTTGTACAAAGTAATGTACTCAAATTGAA<br>GGAAGAAGACAAACAAAAGTTTAATAATGCAGAGCGTTCTGACAAGCGTTGGGTCTT<br>CGTGGACGCGATGAAGAAGAAGCATAGTAAGGTTGGTGAAATGCTCCTTCAGACAT<br>TCTTTAGTGTGGATCCGGGTTCGCATCATGGGGAGGCAAATCTTGAGATGGAAGAA<br>CCGGAGGAGTCATTAAACACGCTCAAACTTTGTTCACCGGAAGAGTTCACCCGCC<br>TTTGCCGGGAGAAGACCCAGGAAATCTATCCGATTAAAGAGGCAAACGGCCGGA<br>CACGTAAGGCCCTTATTATTTGCAACACGGAGTTTAAGCATCTCTCGTTGCGCTATG<br>GTGCCAATTTCGATATTATTGGTATGAAAGGCTTGTTAGAAGACCTTGGGTATGATGTG<br>GTCGTTAAAGAAGAGCTCACTGCCGAGGGGATGGAAAGTGAAATGAAGGATTTCGC<br>AGCCTTATCAGAACACCAAACCTCGGACTCCACGTTTCTTGTACTTATGAGCCACG<br>GCACATTACACGGCATCTGCGGCACAATGCACTCAGAAAAGACACCTGACGTCCT<br>CCAGTACGACACAATCTACCAAATTTTCAATAATTGCCACTGCCCTGGTCTGCGCGA<br>TAAGCCGAAAGTCATCATTGTTCAAGCCGCACGCGGCGGGAATAGCGGCGAAATG<br>TGGATCCGTGAATCTTCGAAGCCACAACTGTGTCGTGGCGTAGATCTTCCGCGTAA<br>CATGGAAGCAGATGCTGTGAAACTCTCGCATGTGGAGAAAGATTTTATTGCCTTTTAT<br>AGTACCACTCCTCACCATTTATCATACCGCGACAAGACCGGTGGTTCGTACTTTATC<br>ACTCGCTTGATCTCTTGCTTTCGCAAACATGCATGTAGCTGTCACCTGTTCGATATTTT<br>CCTGAAAGTTCAGCAATCTTTTGAGAAAGCTAGCATCCACTCCCAAATGCCGACGA<br>TTGACCGCGCTACCTTAACCCGCTACTTTTATTTGTTCCCTGGCAATTAA<br>**Amino Acids:**<br>MKHHHHHHPMSTENLYFQ↓GMAENKHPDKPLKVLEQLGKEVLTEYLEKLVQSNVLK<br>LKEEDKQKFNNAERSDKRWVFVDAMKKKHSKVGEMLLQTFFSVDPGSHHGEANLE<br>MEEPEESLNTLKLCSPEEFTRLCREKTQEIYPIKEANGRTRKALIICNTEFKHLSLRYGA<br>NFDIIGMKGLLEDLGYDVVVKEELTAEGMESEMKDFAALSEHQTSDSTFLVLMSHGTL<br>HGICGTMHSEKTPDVLQYDTIYQIFNNCHCPGLRDKPKVIIVQAARGGNSGEMWIRE<br>SSKPQLCRGVDLPRNMEADAVKLSHVEKDFIAFYSTTPHHLSYRDKTGGSYFITRLISC<br>FRKHACSCHLFDIFLKVQQSFEKASIHSQMPTIDRATLTRYFYLFPGN* | ↓ = TEV cut-site<br>MW = 42,767 Da<br>$\varepsilon_{280}$ = 27,390 $M^{-1}$ $cm^{-1}$ |
| **FL Casp11*** | **DNA:**<br>ATGAAACACCACCACCACCACCACCCGATGAGTACCGAAAATCTGTACTTCCAGG<br>GCGGTGCTGAAAATAAACATCCGGATAAGCCTCTCAAAGTTCTGGAACAATTAGGG<br>AAGGAAGTCCTGACCGAATACCTGGAAAAACTTGTACAAAGTAATGTACTCAAATTGA<br>AGGAAGAAGACAAACAAAAGTTTAATAATGCAGAGCGTTCTGACAAGCGTTGGGTCT<br>TCGTGGACGCGATGAAGAAGAAGCATAGTAAGGTTGGTGAAATGCTCCTTAATAATC<br>TGCTGAGTGTGGATCCGGGTTCGCATCATGGGGAGGCAAATCTTGAGGCAGAAGA<br>ACCGGAGGAGTCAGCAAACAGTGCCAAGAGTGCACCTCACGAAGAGTTCACCCG<br>CGCTTCTCGGGAGAAGACCCAGGAAATCTATCCGATTAAAGAGGCAAACGGCCG<br>GACACGTAAGGCCCTTATTATTTGCAACACGGAGTTTAAGCATCTCTCGTTGCGCTA<br>TGGTGCCAATTTCGATATTATTGGTATGAAAGGCTTGTTAGAAGACCTTGGGTATGATG<br>TGGTCGTTAAAGAAGAGCTCACTGCCGAGGGGATGGAAAGTGAAATGAAGGATTTC<br>GCAGCCTTATCAGAACACCAAACCTCGGACTCCACGTTTCTTGTACTTATGAGCCA | ↓ = TEV cut-site<br>MW = 42,258 Da<br>$\varepsilon_{280}$ = 27,390 $M^{-1}$ $cm^{-1}$ |

| | | |
|---|---|---|
| | CGGCACATTACACGGCATCTGCGGCACATTACACTCAGAAAAGACACCTGACGTC<br>CTCCAGTACGACACAATCTACCAAATTTTCAATAATTGCCACTGCCCTGGTCTGCGC<br>GATAAGCCGAAAGTCATCATTGTTCAAGCCGCACGCGGCGGGAATAGCGGCGAA<br>ATGTGGATCCGTGAATCTTCGAAGCCACAAGCGAGTCGTGGCGTAGATCTTCCGC<br>GTAACCTTGAAGCAGATGCTGTGAAACTCTCGCATGTGGAGAAAGATTTTATTGCCTT<br>TTATAGTACCACTCCTCACCATTTATCATACCGCGACAAGACCGGTGGTTCGTACTTT<br>ATCACTCGCTTGATCTCTTGCTTTCGCAAACATGCATGTAGCTGTCACCTGTTCGATA<br>TTTTCCTGAAAGTTCAGCAATCTTTTGAGAAAGCTAGCATCCACTCCCAATTACCGAC<br>GATTGACCGCGTTACCTTAACCCGCTACTTTTATTTGTTCCCTGGCAATTGA<br>**Amino Acids:**<br>MKHHHHHHPMSTENLYFQG↓GAENKHPDKPLKVLEQLGKEVLTEYLEKLVQSNVLK<br>LKEEDKQKFNNAERSDKRWVFVDAMKKKHSKVGEMLLNNLLSVDPGSHHGEANLE<br>AEEPEESANSAKSAPHEEFTRASREKTQEIYPIKEANGRTRKALIICNTEFKHLSLRYGA<br>NFDIIGMKGLLEDLGYDVVVKEELTAEGMESEMKDFAALSEHQTSDSTFLVLMSHGTL<br>HGICGTLHSEKTPDVLQYDTIYQIFNNCHCPGLRDKPKVIIVQAARGGNSGEMWIRES<br>SKPQASRGVDLPRNLEADAVKLSHVEKDFIAFYSTTPHHLSYRDKTGGSYFITRLISCF<br>RKHACSCHLFDIFLKVQQSFEKASIHSQLPTIDRVTLTRYFYLFPGN* | |
| **CARD-TrpCage** | **DNA:**<br>ATGGCTGAAAATAAACATCCGGATAAGCCTCTCAAAGTTCTGGAACAATTAGGGAAG<br>GAAGTCCTGACCGAATACCTGGAAAAACTTGTACAAAGTAATGTACTCAAATTGAAG<br>GAAGAAGACAAACAAAAGTTTAATAATGCAGAGCGTTCTGACAAGCGTTGGGTCTTC<br>GTGGACGCGATGAAGAAGAAGCATAGTAAGGTTGGTGAAATGCTCCTTCAGACATT<br>CTTTAGTGTGGATCCGGGTTCGCATCATGGGGAGGCAAATCTTGAGATGGAAGAAC<br>CGGAGGAGTCATTAAACACGCTCAAACTTTGTTCACCGGAAGAGTTCACCCGCCTT<br>TGCCGGGAGGGCGATGCGTATGCCCAGTGGTTGAAAGATGGGGGTCCGTCATCT<br>GGCCGCCCGCCGCCGTCATCGGGCTCGCATCACCACCACCATCACTAA<br>**Amino Acids:**<br>MAENKHPDKPLKVLEQLGKEVLTEYLEKLVQSNVLKLKEEDKQKFNNAERSDKRWVF<br>VDAMKKKHSKVGEMLLQTFFSVDPGSHHGEANLEMEEPEESLNTLKLCSPEEFTRLC<br>REGDAYAQWLKDGGPSSGRPPPSSGSHHHHHH* | MW = 16,578 Da<br>$\varepsilon_{280}$ = 13,980 $M^{-1}$ $cm^{-1}$<br>= 0.843 L $g^{-1}$ $cm^{-1}$<br>dn/dc = 0.187 mL/g |
| **CARD-GB1** | **DNA:**<br>ATGGCTGAAAATAAACATCCGGATAAGCCTCTCAAAGTTCTGGAACAATTAGGGAAG<br>GAAGTCCTGACCGAATACCTGGAAAAACTTGTACAAAGTAATGTACTCAAATTGAAG<br>GAAGAAGACAAACAAAAGTTTAATAATGCAGAGCGTTCTGACAAGCGTTGGGTCTTC<br>GTGGACGCGATGAAGAAGAAGCATAGTAAGGTTGGTGAAATGCTCCTTCAGACATT<br>CTTTAGTGTGGATCCGGGTTCGCATCATGGGGAGGCAAATCTTGAGATGGAAGAAC<br>CGGAGGAGTCATTAAACACGCTCAAACTTTGTTCACCGGAAGAGTTCACCCGCCTT<br>TGCCGGGAGAGTTACAAACTCATTCTGAATGGCAAGACATTAAAGGGTGAAACAAC<br>TACTGAGGCAGTGGATGCCGCCACGGCAGAAAAAGTTTTCAAACAGTATGCCAATG<br>ACAATGGTGTGGATGGTGAGTGGACCTATGATGACGCGACGAAAACGTTCACGGTA<br>ACCGAATCGGGCTCGCATCACCACCACCATCACTAA<br>**Amino Acids:**<br>MAENKHPDKPLKVLEQLGKEVLTEYLEKLVQSNVLKLKEEDKQKFNNAERSDKRWVF<br>VDAMKKKHSKVGEMLLQTFFSVDPGSHHGEANLEMEEPEESLNTLKLCSPEEFTRLC<br>RESYKLILNGKTLKGETTTEAVDAATAEKVFKQYANDNGVDGEWTYDDATKTFTVTESG<br>SHHHHHH* | MW = 20,485 Da<br>$\varepsilon_{280}$ = 16,960 $M^{-1}$ $cm^{-1}$<br>= 0.828 L $g^{-1}$ $cm^{-1}$<br>dn/dc = 0.188 mL/g |
| **CARD-SUMO** | **DNA:**<br>ATGGCTGAAAATAAACATCCGGATAAGCCTCTCAAAGTTCTGGAACAATTAGGGAAG<br>GAAGTCCTGACCGAATACCTGGAAAAACTTGTACAAAGTAATGTACTCAAATTGAAG<br>GAAGAAGACAAACAAAAGTTTAATAATGCAGAGCGTTCTGACAAGCGTTGGGTCTTC<br>GTGGACGCGATGAAGAAGAAGCATAGTAAGGTTGGTGAAATGCTCCTTCAGACATT<br>CTTTAGTGTGGATCCGGGTTCGCATCATGGGGAGGCAAATCTTGAGATGGAAGAAC<br>CGGAGGAGTCATTAAACACGCTCAAACTTTGTTCACCGGAAGAGTTCACCCGCCTT<br>TGCCGGGAGAGCGGCCTGGTTCCGCGTGGGAGTGCAAGCATGTCGGACAGCGA<br>AGTTAATCAAGAAGCCAAACCGGAGGTCAAACCGGAAGTTAAGCCAGAGACTCATA<br>TCAACTTAAAAGTTTCAGATGGGAGTAGCGAGATCTTTTTTAAGATCAAAAAAACAAC<br>CCCGCTGCGCCGTCTGATGGAAGCATTTGCGAAACGCCAAGGCAAAGAAATGGA<br>CTCCTTGCGTTTTCTGTATGATGGGATTCGCATTCAAGCGGATCAGACGCCGGAAG | MW = 26,494 Da<br>$\varepsilon_{280}$ = 8,480 $M^{-1}$ $cm^{-1}$<br>= 0.320 L $g^{-1}$ $cm^{-1}$<br>dn/dc = 0.187 mL/g |

| | | |
|---|---|---|
| | ACCTTGACATGGAGGATAACGATATTATCGAAGCTCACCGCGAACAAATTTCGGGC<br>TCGCATCACCACCACCATCACTAA<br>**Amino Acids:**<br>MAENKHPDKPLKVLEQLGKEVLTEYLEKLVQSNVLKLKEEDKQKFNNAERSDKRWVF<br>VDAMKKKHSKVGEMLLQTFFSVDPGSHHGEANLEMEEPEESLNTLKLCSPEEFTRLC<br>RESGLVPRGSASMSDSEVNQEAKPEVKPEVKPETHINLKVSDGSSEIFFKIKKTTPLRR<br>LMEAFAKRQGKEMDSLRFLYDGIRIQADQTPEDLDMEDNDIIEAHREQISGSHHHHH<br>H* | |
| **WT Casp11 PD** | **DNA:**<br>*His6MBPtag*...CTCGGGGAAAATTTATACTTCCAGGGCATGGAAGAACCGGAGGAG<br>TCATTAAACACGCTCAAACTTTGTTCACCGGAAGAGTTCACCCGCCTTTGCCGGGA<br>GAAGACCCAGGAAATCTATCCGATTAAAGAGGCAAACGGCCGGACACGTAAGGC<br>CCTTATTATTTGCAACACGGAGTTTAAGCATCTCTCGTTGCGCTATGGTGCCAATTTC<br>GATATTATTGGTATGAAAGGCTTGTTAGAAGACCTTGGGTATGATGTGGTCGTTAAAGA<br>AGAGCTCACTGCCGAGGGGATGGAAAGTGAAATGAAGGATTTCGCAGCCTTATCA<br>GAACACCAAACCTCGGACTCCACGTTTCTTGTACTTATGAGCCACGGCACATTACA<br>TGGCATCTGCGGCACAATGCACTCAGAAAAGACACCTGACGTCCTCCAGTACGAC<br>ACAATCTACCAAATTTTCAATAATTGCCACTGCCCTGGTCTGCGCGATAAGCCGAAA<br>GTCATCATTGTTCAAGCCTGCCGCGGCGGGAATAGCGGCGAAATGTGGATCCGTG<br>AATCTTCGAAGCCACAACTGTGTCGTGGCGTAGATCTTCCGCGTAACATGGAAGCA<br>GATGCTGTGAAACTCTCGCATGTGGAGAAAGATTTTATTGCCTTTTATAGTACCACTC<br>CTCACCATTTATCATACCGCGACAAGACCGGTGGTTCGTACTTTATCACTCGCTTGA<br>TCTCTTGCTTTCGCAAACATGCATGTAGCTGTCACCTGTTCGATATTTTCCTGAAAGTT<br>CAGCAATCTTTTGAGAAAGCTAGCATCCACTCCCAAATGCCGACGATTGACCGCG<br>CTACCTTAACCCGCTACTTTTATTTGTTCCCTGGCAATTAA<br>**Amino Acids:**<br>*His6MBPtag*...LGENLYFQ↓GMEEPEESLNTLKLCSPEEFTRLCREKTQEIYPIKEANG<br>RTRKALIICNTEFKHLSLRYGANFDIIGMKGLLEDLGYDVVVKEELTAEGMESEMKDFA<br>ALSEHQTSDSTFLVLMSHGTLHGICGTMHSEKTPDVLQYDTIYQIFNNCHCPGLRDK<br>PKVIIVQACRGGNSGEMWIRESSKPQLCRGVDLPRNMEADAVKLSHVEKDFIAFYSTT<br>PHHLSYRDKTGGSYFITRLISCFRKHACSCHLFDIFLKVQQSFEKASIHSQMPTIDRAT<br>LTRYFYLFPGN* | ↓ = TEV cut-site<br>MW = 32,254 Da<br>$\varepsilon_{280}$ = 20,400 $M^{-1}$ $cm^{-1}$ |

*The MW, $\varepsilon_{280}$, and dn/dc values are calculated based on the underlined amino sequence

**Table S2.** Sample and buffer conditions for experiments in this study.

| Method/Samples | SEC/Analysis Buffer components* | Location |
|---|---|---|
| **NMR** | | |
| 125 μM Casp11 CARD [U-$^{13}$C,$^{15}$N] | 20 mM NaPhos, 100 mM NaCl, 1 mM EDTA, 3% $D_2O$, pH 6.0 | Fig. 1B(left), B(i), C<br>Fig. S2A(left) |
| 120 μM Casp11 CARD [U-$^{15}$N] | 20 mM NaPhos, 100 mM NaCl, 1 mM EDTA, 3% $D_2O$, pH 6.0 | Fig. 1B(ii)<br>Fig.S2B |
| 105 μM Casp11 CARD [U-$^{13}$C,$^{15}$N] | **SEC:** 20 mM NaPhos, 100 mM NaCl, 1 mM EDTA, 3% $D_2O$, pH 6.0<br>**Analysis:** 0-6 M Urea, 20 mM NaPhos, 100 mM NaCl, 1 mM EDTA, 3% $D_2O$, pH 6.0 | Fig. S3B-C |
| 720 μM Casp11 CARD [U-$^{13}$C,$^{15}$N]‡ | **Analysis:** 6 M Urea, 20 mM NaPhos, 100 mM NaCl, 1 mM EDTA, 3% $D_2O$, pH 6.0 | Fig. S2A(right), B<br>Fig. S3A |
| 85 μM Casp11 CARD [U-$^2$H; MLV] | 20 mM NaPhos, 100 mM NaCl, 1 mM EDTA, 3% $D_2O$, pH 7.4 | Fig. 2B |
| 75 μM Casp11 CARD [U-$^2$H; MLV] + 1×KLA [U-$^1$H] (SEC-filtered) | | |
| 80 μM Casp11 CARD-MMM [U-$^1$H; MLV] + 3×KLA [U-$^1$H] (SEC-filtered) | **SEC:** 20 mM Bicine, 150 mM NaCl, 15 mM TCEP, 1 mM EDTA, 3% $D_2O$, pH 8.0<br>**Analysis:** 20 mM Bicine, 50 mM NaCl, 15 mM TCEP, 1 mM EDTA, 3% $D_2O$, pH 8.0 | Fig. 3B |
| 50 μM C254A FL Casp11$^{PR}$ [U-$^1$H; MLV] + 3×KLA [U-$^1$H] (SEC-filtered) | | |
| 10-300 μM Casp11* PD [U-$^1$H; IM] | **Analysis:** 20 mM Bicine, 50 mM NaCl, 15 mM TCEP, 1 mM EDTA, 99.9% $D_2O$, pD 8.0 | Fig. 3C |
| 10-60 μM FL Casp11* [U-$^1$H; IM] + 3×KLA [U-$^1$H] (SEC-filtered) | **SEC:** 20 mM Bicine, 150 mM NaCl, 15 mM TCEP, 1 mM EDTA, 99.9% $D_2O$, pD 8.0<br>**Analysis:** 20 mM Bicine, 50 mM NaCl, 15 mM TCEP, 1 mM EDTA, 99.9% $D_2O$, pD 8.0 | Fig. 3D |
| 350 & 830§ μM C254 D277A D285A Casp11 PD [U-$^1$H; IM] | **Analysis:** 20 mM Bicine, 50 mM NaCl, 15 mM TCEP, 1 mM EDTA, 3% $D_2O$, pH 8.0 | Fig. S8A |
| 360 μM C254A Casp11 PD [U-$^2$H; ILVM] | **Analysis:** 20 mM HEPES, 15 mM TCEP, 0.5 mM EDTA, 99.9% $D_2O$, pD 8.0 | Fig. S9 |
| 600 μM Casp11* PD [U-$^1$H; IM] | **Analysis:** 20 mM Bicine, 50 mM NaCl, 15 mM TCEP, 1 mM EDTA, 99.9% $D_2O$, pD 8.0 | |
| 30 μM Casp11 CARD-His6 [U-$^{15}$N] (prepared under denaturing conditions) | 20 mM NaPhos, 100 mM NaCl, 1 mM EDTA, 3% $D_2O$, pH 6.0 | Inset in *SI Appendix*, '*Casp11 expression and purification*' |
| 30 μM Casp11 CARD-His6 [U-$^{15}$N] (prepared under native buffer conditions) | | |
| ‡ - Indicates SEC step was skipped. Instead buffer exchanged (>1000×) directly into analysis buffer.<br>§ - Indicates a 3mm NMR tube was used. In all other cases a 5mm NMR tube was used. | | |
| **CD** | | |
| 30 μM Casp11 CARD | **SEC:** 10 mM NaPhos, 150 mM NaF, 0.5 mM EDTA, pH 7.4<br>**Analysis:** 10 mM NaPhos, 90 mM NaF, 0.5 mM EDTA, pH 7.4 | Fig. S4 |
| 30 μM Casp11 CARD + 3×KLA | | |
| **SEC-UV-RI-LS** | | |
| 110 & 115 μM Casp11 CARD + 3×KLA (SEC-filtered) | 20 mM NaPhos, 150 mM NaCl, 1 mM EDTA, pH 7.4 | Fig. 2A |

| | | |
|---|---|---|
| 110 & 160 μM CARD-TrpCage + 3×KLA (SEC-filtered) | 20 mM NaPhos, 150 mM NaCl, 1 mM EDTA, pH 7.4 | Fig. S5 |
| 80 & 110 μM CARD-GB1 + 3×KLA (SEC-filtered) | | |
| 105 & 145 μM CARD-SUMO + 3×KLA (SEC-filtered) | | |
| **SEC-UV** | | |
| 35 & 50 μM C254A FL Casp11 + 0-5×KLA | 20 mM NaPhos, 150 mM NaCl, 1 mM EDTA, pH 7.4 | Fig. 3A<br>Fig. S7 (3×KLA only) |
| 50 μM C254A FL Casp11$^{PR}$ + 3×KLA | | Fig. S7 |
| **Cryo-EM** | | |
| 85 μM C254A FL Casp11 + 3×KLA (SEC-filtered) | **SEC:** 20 mM Bicine, 50 mM NaCl, 10 mM TCEP, 1 mM EDTA, pH 8.0<br>**Analysis:** 20 mM Bicine, 50 mM NaCl, 10 mM TCEP, 1 mM EDTA, pH 8.0 | Fig. S6 |
| **AUC** | | |
| 0.1-19 μM WT Casp11 PD | **Analysis:** 10 mM Tris, 0.5 mM TCEP, 80 mM NaCl, pH 8.0 | Fig. S8B |

*If not specified, buffer conditions for SEC and analysis are identical

### References


1. B. Dang, *et al.*, SNAC-tag for sequence-specific chemical protein cleavage. *Nat. Methods* **16**, 319–322 (2019).

2. T. G. M. Schmidt, *et al.*, Development of the Twin-Strep-tag® and its application for purification of recombinant proteins from cell culture supernatants. *Protein Expr. Purif.* **92**, 54–61 (2013).

3. P. J. Shilling, *et al.*, Improved designs for pET expression plasmids increase protein production yield in Escherichia coli. *Commun. Biol.* **3**, 214 (2020).

4. L. Kay, P. Keifer, T. Saarinen, Pure absorption gradient enhanced heteronuclear single quantum correlation spectroscopy with improved sensitivity. *J. Am. Chem. Soc.* **114**, 10663–10665 (1992).

5. J. An, *et al.*, Caspase-4 disaggregates lipopolysaccharide micelles via LPS-CARD interaction. *Sci. Rep.* **9**, 826 (2019).

6. K. H. Sumida, *et al.*, Improving Protein Expression, Stability, and Function with ProteinMPNN. *J. Am. Chem. Soc.* **146**, 2054–2061 (2024).

7. C. R. H. Raetz, *et al.*, Kdo2-Lipid A of Escherichia coli, a defined endotoxin that activates macrophages via TLR-4. *J. Lipid Res.* **47**, 1097–1111 (2006).

8. F. Delaglio, *et al.*, NMRPipe: A multidimensional spectral processing system based on UNIX pipes. *J. Biomol. NMR* **6** (1995).

9. J. J. Helmus, C. P. Jaroniec, Nmrglue: an open source Python package for the analysis of multidimensional NMR data. *J. Biomol. NMR* **55**, 355–367 (2013).

10. M. Sattler, C. Griesinger, J. Schleucher, Heteronuclear multidimensional NMR experiments for the structure determination of proteins in solution employing pulsed field gradients. *Prog. Nucl. Magn. Reson. Spectrosc.* **34**, 93–158 (1999).

11. S. C. Panchal, N. S. Bhavesh, R. V. Hosur, Improved 3D triple resonance experiments, HNN and HN(C)N, for $H^N$ and $^{15}N$ sequential correlations in ($^{13}C$, $^{15}N$) labeled proteins: Application to unfolded proteins. *J. Biomol. NMR* **20**, 135–147 (2001).

12. M. Ikura, L. E. Kay, A. Bax, A novel approach for sequential assignment of proton, carbon-13, and nitrogen-15 spectra of larger proteins: heteronuclear triple-resonance three-dimensional NMR spectroscopy. Application to calmodulin. *Biochemistry* **29**, 4659–4667 (1990).

13. S. G. Hyberts, K. Takeuchi, G. Wagner, Poisson-Gap Sampling and Forward Maximum Entropy Reconstruction for Enhancing the Resolution and Sensitivity of Protein NMR Data. *J. Am. Chem. Soc.* **132**, 2145–2147 (2010).

14. J. Ying, F. Delaglio, D. A. Torchia, A. Bax, Sparse multidimensional iterative lineshape-enhanced (SMILE) reconstruction of both non-uniformly sampled and conventional NMR data. *J. Biomol. NMR* **68**, 101–118 (2017).

15. W. Lee, M. Tonelli, J. L. Markley, NMRFAM-SPARKY: enhanced software for biomolecular NMR spectroscopy. *Bioinformatics* **31**, 1325–1327 (2015).

16. W. Lee, *et al.*, I-PINE web server: an integrative probabilistic NMR assignment system for proteins. *J. Biomol. NMR* **73**, 213–222 (2019).

17. D. F. Hansen, P. Vallurupalli, L. E. Kay, An Improved $^{15}$N Relaxation Dispersion Experiment for the Measurement of Millisecond Time-Scale Dynamics in Proteins. *J. Phys. Chem. B* **112**, 5898–5904 (2008).

18. W.-Y. Choy, *et al.*, Distribution of molecular size within an unfolded state ensemble using small-angle X-ray scattering and pulse field gradient NMR techniques. *J. Mol. Biol.* **316**, 101–112 (2002).

19. R. Huang, F. Pérez, L. E. Kay, Probing the cooperativity of *Thermoplasma acidophilum* proteasome core particle gating by NMR spectroscopy. *Proc. Natl. Acad. Sci.* **114**, E9846–E9854 (2017).

20. G. A. Morris, R. Freeman, Enhancement of nuclear magnetic resonance signals by polarization transfer. *J. Am. Chem. Soc.* **101**, 760–762 (1979).

21. K. Wang, *et al.*, Structural Mechanism for GSDMD Targeting by Autoprocessed Caspases in Pyroptosis. *Cell* **180**, 941-955.e20 (2020).

22. Y. Zhou, Q. Pan, D. E. V. Pires, C. H. M. Rodrigues, D. B. Ascher, DDMut: predicting effects of mutations on protein stability using deep learning. *Nucleic Acids Res.* **51**, W122–W128 (2023).

23. D. J. Slotboom, R. H. Duurkens, K. Olieman, G. B. Erkens, Static light scattering to characterize membrane proteins in detergent solution. *Methods* **46**, 73–82 (2008).

24. H. Zhao, P. H. Brown, P. Schuck, On the Distribution of Protein Refractive Index Increments. *Biophys. J.* **100**, 2309–2317 (2011).

25. A. Theisen, Ed., *Refractive increment data-book for polymer and biomolecular scientists* (Nottingham University Press, 2000).

26. T. Tumolo, L. Angnes, M. S. Baptista, Determination of the refractive index increment (dn/dc) of molecule and macromolecule solutions by surface plasmon resonance. *Anal. Biochem.* **333**, 273–279 (2004).

27. Eli. Grushka, Characterization of exponentially modified Gaussian peaks in chromatography. *Anal. Chem.* **44**, 1733–1738 (1972).

28. Y. Kalambet, Y. Kozmin, K. Mikhailova, I. Nagaev, P. Tikhonov, Reconstruction of chromatographic peaks using the exponentially modified Gaussian function. *J. Chemom.* **25**, 352–356 (2011).

29. A. Micsonai, *et al.*, Accurate secondary structure prediction and fold recognition for circular dichroism spectroscopy. *Proc. Natl. Acad. Sci.* **112** (2015).

30. A. Micsonai, *et al.*, BeStSel: a web server for accurate protein secondary structure prediction and fold recognition from the circular dichroism spectra. *Nucleic Acids Res.* **46**, W315–W322 (2018).

31. N. Sreerama, S. Yu. Venyaminov, R. W. Woody, Estimation of Protein Secondary Structure from Circular Dichroism Spectra: Inclusion of Denatured Proteins with Native Proteins in the Analysis. *Anal. Biochem.* **287**, 243–251 (2000).

32. G. Nagy, H. Grubmuller, Implementation of a Bayesian secondary structure estimation method for the SESCA circular dichroism analysis package. *Comput. Phys. Commun.* **266**, 108022 (2021).

33. O. Burastero, *et al.*, ChiraKit: an online tool for the analysis of circular dichroism spectroscopy data. *Nucleic Acids Res.* **53**, W158–W168 (2025).

34. C. R. Marr, S. Benlekbir, J. L. Rubinstein, Fabrication of carbon films with ~500nm holes for cryo-EM with a direct detector device. *J. Struct. Biol.* **185**, 42–47 (2014).

35. H. Guo, *et al.*, Electron-event representation data enable efficient cryoEM file storage with full preservation of spatial and temporal resolution. *IUCrJ* **7**, 860–869 (2020).

36. A. Punjani, J. L. Rubinstein, D. J. Fleet, M. A. Brubaker, cryoSPARC: algorithms for rapid unsupervised cryo-EM structure determination. *Nat. Methods* **14**, 290–296 (2017).

37. H. Zhao, *et al.*, Quantitative Analysis of Protein Self-Association by Sedimentation Velocity. *Curr. Protoc. Protein Sci.* **101**, e109 (2020).

38. P. Schuck, Size-Distribution Analysis of Macromolecules by Sedimentation Velocity Ultracentrifugation and Lamm Equation Modeling. *Biophys. J.* **78**, 1606–1619 (2000).

39. C. A. Brautigam, "Calculations and Publication-Quality Illustrations for Analytical Ultracentrifugation Data" in *Methods in Enzymology*, (Elsevier, 2015), pp. 109–133.

40. J. S. Philo, SEDNTERP: a calculation and database utility to aid interpretation of analytical ultracentrifugation and light scattering data. *Eur. Biophys. J.* **52**, 233–266 (2023).

41. K. P. Downs, H. Nguyen, A. Dorfleutner, C. Stehlik, An overview of the non-canonical inflammasome. *Mol. Aspects Med.* **76**, 100924 (2020).

42. N. Kayagaki, *et al.*, Non-canonical inflammasome activation targets caspase-11. *Nature* **479**, 117–121 (2011).

43. V. A. K. Rathinam, *et al.*, TRIF licenses caspase-11-dependent NLRP3 inflammasome activation by gram-negative bacteria. *Cell* **150**, 606–619 (2012).

44. S. Rühl, P. Broz, Caspase-11 activates a canonical NLRP3 inflammasome by promoting K(+) efflux. *Eur. J. Immunol.* **45**, 2927–2936 (2015).

45. T. D. Goddard, *et al.*, UCSF ChimeraX: Meeting modern challenges in visualization and analysis: UCSF ChimeraX Visualization System. *Protein Sci.* **27**, 14–25 (2018).

46. J. Jumper, *et al.*, Highly accurate protein structure prediction with AlphaFold. *Nature* **596**, 583–589 (2021).

47. M. Varadi, *et al.*, AlphaFold Protein Structure Database: massively expanding the structural coverage of protein-sequence space with high-accuracy models. *Nucleic Acids Res.* **50**, D439–D444 (2022).
